# Variations in carbapenem resistance associated with the VIM-1 metallo-β-lactamase across the *Enterobacterales*

**DOI:** 10.1101/2025.09.07.674714

**Authors:** Mia Rondinelli, Sabhjeet Kaur, Owen A. Ledwell, Henry Wong, Prameet M. Sheth, George C. diCenzo

## Abstract

The VIM-1 metallo-β-lactamase enzyme, encoded as a cassette within class 1 integrons, is found in Gram-negative clinical isolates worldwide and has been linked to outbreaks of bacterial pathogens in nosocomial settings. Six *vim-1^+^* clinical isolates, from the genera *Escherichia*, *Klebsiella*, and *Enterobacter*, were obtained from Kingston, Ontario, Canada. Whole genome sequencing revealed that *vim-1* was plasmid-borne in all strains and situated as the first gene in In916 or In110 integrons. Analysis of related plasmids suggested that these *vim-1*-containing plasmids are globally disseminated and have spread via horizontal gene transfer and autochthonous vertical spread within Ontario. Interestingly, the minimum inhibitory concentrations of ertapenem and meropenem, two clinically relevant carbapenem antibiotics, against these six isolates varied more than tenfold, suggesting the effects of VIM-1 are dependent on the genomic content of the host microbe. To further study the genomic content dependency of VIM-1, we introduced *vim-1* into three common *Enterobacterales* laboratory strains. Although introduction of *vim-1* into *Escherichia coli* DH5α resulted in little resistance to ertapenem or meropenem, multiple rounds of adaptive laboratory evolution allowed us to identify variants with extremely high levels of resistance to both carbapenems. DNA sequencing revealed that the increase in carbapenem resistance was due to a combination of increased *vim-1* gene dosage and epistatic interactions with mutations of *ompC* that likely would have decreased outer membrane permeability to these antibiotics. Together, these results provide additional support for the role of gene epistasis is modulating the antimicrobial resistance phenotypes of acquired resistance genes, as well as previous results suggesting that the presence of a β-lactamase gene is insufficient to confer strong resistance to carbapenems without being paired with reduced outer membrane permeability.

## INTRODUCTION

Antimicrobial resistant pathogens are three times more likely to result in mortality than their susceptible counterparts (Dadgostar, 2019), resulting in an estimated global death toll of 1.27 million in 2019 alone (Murray et al., 2022). In bacteria, horizontal gene transfer (HGT) facilitates the exchange of antimicrobial resistance genes (ARGs) across diverse bacterial populations and the collection of these genes into conjugative transposons (van der Zee et al., 2018) and plasmids (De la Cruz Barrón et al., 2018). These, in turn, disseminate quickly across microbial ecosystems via HGT (von Wintersdorff et al., 2016), occasionally recombining with each other along the way (Y. Li et al., 2018). Transposons can additionally exacerbate antimicrobial resistance (AMR) by inserting into promoter regions, activating genes associated with conjugation, and improving overall conjugation frequency (Poidevin et al., 2018).

While HGT plays a role in ARG dissemination and evolution, there appear to be limits to how far ARGs can spread, suggesting interplay between the chromosomal genome and acquired genes (Wong, 2017). Studies investigating the dissemination of resistance to antimicrobial peptides (AMPs) and antibiotics have noted that the transfer of ARGs occurs at a higher rate between more closely related bacteria (Hu et al., 2016; Kintses et al., 2019). In one case, genes encoding AMPs from gut microbiota could not confer resistance when transferred to *Escherichia coli* (Kintses et al., 2019). In others, the transfer of ARGs from distantly related species (such as *Shewanella* spp.) to *E. coli* resulted in toxicity and cell death (Sorek et al., 2007). This is likely due to the dependence of resistance mechanisms on host physiology, suggesting that there are phylogenetic barriers that may limit the spread of certain ARGs (Porse et al., 2018; Sorek et al., 2007).

Additionally, there are examples of compensatory mutations increasing the fitness of bacterial strains that have evolved AMR mechanisms. Knopp and Andersson (2015) found that *E. coli* could overcome the fitness cost of losing outer membrane proteins C and F through compensatory mutations to pathways producing other porins. There is also evidence that epistatic relationships exist between acquired and chromosomal ARGs. For instance, Silva et al (2011) found that, occasionally, plasmids carrying resistance genes confer a selective advantage to strains carrying ARGs on their chromosome in the absence of antibiotics.

Surveillance data from across Southern Ontario identified the emergence of clinical isolates in Gram-negative *Enterobacterales* carrying the Verona Integron-encoded Metallo-β-lactamase (VIM) (P. P. Kohler et al., 2018) that was first identified in Europe (Arcari et al., 2020; P. Kohler et al., 2020; Mano et al., 2015; Papadimitriou-Olivgeris et al., 2019). Unlike other metallo-β-lactamases (MBLs) identified in this Canadian surveillance program (Kohler et al., 2020), VIM-positive patients had none of the ‘classical’ risk factors associated seen in patients with other MBL’s including documented travel history outside of Canada or hospital admission, suggesting local acquisition. The dissemination of these pathogens is likely through undetected colonization and transmission amongst patients in acute care (Kohler et al., 2020) or acquisition via unknown community exposure.

Like other MBLs, but unlike extended-spectrum β-lactamases (ESBLs), VIM can hydrolyze carbapenem antibiotics, which are often considered as a last-resort antibiotic. Of all VIM genes, *vim-1* is the most common gene variant globally (Matsumura et al., 2017). Like other B1 MBLs, the structure of VIM-1 displays an overall αβ/βα fold with the Zn^2+^ centre forming the active site situated in a shallow groove formed by the interface of the two αβ domains (Salimraj et al., 2019). The *vim-1* gene is generally carried as part of class 1 integrons, which are major contributors to AMR through their ability to capture and express a diverse range of ARGs (Gillings et al., 2008). Class 1 integrons can be found embedded within an extremely broad range of plasmids, including plasmids of incompatibility (Inc) groups IncA (Botelho et al., 2018), IncC (Gaibani et al., 2018), IncH12 (Kohler et al., 2020), IncFII (Sánchez-Romero et al., 2012), IncN (Miriagou et al., 2010), IncP (Zeng et al., 2019), IncL/M (Peirano et al., 2014), and IncR (P. Kohler et al., 2020). There is also evidence of these integrons integrating into the host chromosome (Irrgang et al., 2019). This promiscuity results in highly mobile genetic apparatuses that can spread horizontally across a broad range of bacterial species.

Here, we report the sequencing and phenotypic characterization of six *Enterobacterales* clinical isolates collected at Kingston Health Sciences Center, all of which were positive for *vim-1.* We demonstrate that the minimum inhibitory concentrations (MIC) of the carbapenem antibiotics ertapenem and meropenem varied more than tenfold across the isolates, suggesting that the effects of *vim-1* are dependent on the genomic content of the host microbe. Introduction of *vim-1* into common laboratory strains resulted in little ertapenem resistance. However, adaptive laboratory evolution led to massive increases in the level of *vim-1*-mediated ertapenem resistance, which we show was due to a combination of increased *vim-1* gene dosage and epistatic interactions with mutations of the *ompC* gene encoding an outer membrane protein.

## MATERIALS AND METHODS

### Bacterial strains and growth conditions

Bacterial strains used in this work are listed in **Table 1**. Strains were routinely grown using Lysogeny Broth (LB) medium (10 g L^−1^ tryptone, 5 g L^−1^ yeast extract, 5 g L^−1^ sodium chloride) and Mueller Hinton (MH) medium (Sigma Aldrich; Product No. 90922). Gentamicin (10 µg/mL in liquid media, 20 µg/mL in solid media) was used to maintain pBBR1mcs-5 plasmids in transformed strains. Super optimal broth with catabolite repression (SOC) (20 g L^−1^ tryptone, 5 g L^−1^ yeast extract, 10 mM NaCl, 2.5 mM KCl, 10 mM MgSO4, 10 mM CaCl2, 20 mM glucose) was used as a recovery medium following transformation by electroporation. All strains were grown at 37°C. When required, 5-bromo-4-chloro-3-indolyl-β-D-galactopyranoside (X-gal) was added to the media at a final concentration of 40 µg mL^−1^.

**Table 1.** Bacterial strains and plasmids.

| Strain/Plasmid | Species | Characteristics | Year Isolated | Source | Reference |
| --- | --- | --- | --- | --- | --- |
| DH5α | *Escherichia coli* | F<sup>-</sup> *endA1 glnV44 thi-1 recA1 relA1 gyrA96 deoR nupG purB20*<br>φ80dlacZΔM15 Δ(*lacZYA-argF*)U169, hsdR17(*rK<sup>-</sup>mK<sup>+</sup>*), λ<sup>-</sup> | n/a | n/a | Grant et al. (1990) |
| MG1655 | *Escherichia coli* | F<sup>-</sup> λ<sup>-</sup> *ilvG<sup>-</sup> rfb-50 rph-1* | n/a | n/a | Guyer et al. (1981) |
| F5446 | *Escherichia coli* | Clinical isolate from KHSC | 2015 | Blood & urine | This study |
| S2568 | *Escherichia coli* | Clinical isolate from KHSC | 2019 | Rectal screen & urine | This study |
| F48994 | *Enterobacter hormaechei* | Clinical isolate from KHSC | 2018 | Urine | This study |
| H17629 | *Enterobacter hormaechei* | Clinical isolate from KHSC | 2002 | Wound swab | This study |
| H70375 | *Enterobacter hormaechei* | Clinical isolate from KHSC | 2015 | Blood culture | This study |
| T64870 | *Klebsiella pneumoniae* | Clinical isolate from KHSC | 2017 | Sputum culture | This study |
| KP5022 | *Klebsiella grimontii* | hisD2 hsdRI nif<sup>+</sup> | n/a | n/a | Liu et al. (2018) |
| pBBRmcs-5 | - | Broad host range vector; pLac promoter; RK2 mobilizable; Gm<sup>R</sup> | n/a | n/a | Kovach et al. (1995) |

### Whole genome sequencing, assembly, and annotation

Genomic DNA (gDNA) samples were isolated from clinical isolates using phenol-chloroform extraction (Cowie et al., 2006). Purified gDNA samples were then sequenced using an Oxford Nanopore Technologies (ONT) MinION Mk1B nanopore sequencer and the Rapid Barcoding Kit (RBK004) according to the manufacturer’s instructions. Basecalling and demultiplexing were performed using GPU-enabled Guppy version 5.011+2b6dbffa5 and the High-Accuracy model (Oxford Nanopore Technologies). Purified gDNA samples were also sequenced at the Microbial Genome Sequencing Center (Pittsburgh, PA, USA) using an Illumina NextSeq 550 instrument with 150 bp paired-end reads.

Genome assemblies were generated using a pipeline previously described by Duan et al (2022). Programs and versions used are as follows: Flye version 2.8.3 (Kolmogorov et al., 2019), NUCmer version 4.0.0rc1 (Kurtz et al., 2004), Racon version 1.4.22 (Vaser et al., 2017), Minimap2 version 2.20-r1061 (Li, 2018), Medaka version 1.4.1 (Oxford Nanopore Technologies), Pilon version 1.24 (Walker et al., 2014), bwa version 0.7.17.r1198-dirty (H. Li & Durbin, 2009), and Circlator version 1.5.5 (Hunt et al., 2015). Assembly quality was checked using CheckM version 1.2.2 (Parks et al., 2015).

Genome assemblies were annotated using the NCBI prokaryotic genome annotation pipeline (PGAP) build 5429 (Tatusova et al., 2016). Plasmid replicons were identified in the genome assemblies using PlasmidFinder version 2.1.1 (Carattoli et al., 2014). Integron-associated features were identified within the genome assemblies using IntegronFinder version 2.0.2 (Néron et al., 2022). Resistance genes/proteins were identified in nucleotide or amino acid sequences using the Comprehensive Antibiotic Resistance Database (CARD) 3.2.2 Resistance Gene Identifier (RGI) 5.2.1 (Alcock et al., 2020). Genome assemblies were visualized in Mauve snapshot 2015-02-25 build 0 following alignment with progressiveMauve (Darling et al., 2004). Taxonomic classification of strains was performed using TYGS (Meier-Kolthoff et al., 2022).

### Identification of related *vim-1*-containing plasmids

To identify *vim-1*-containing plasmids related to those of our clinical isolates, we first downloaded the plasmid database (PLSDB) version 2021_06_23_v2 (Galata et al., 2019). Next, each of the *vim-1*-containing plasmids of our clinical isolates were individually used as queries with BLASTn from BLAST version 2.10.1+ (Camacho et al., 2009) to search PLSDB, and the top 20 hits for each plasmid were recorded. The exception was the *vim-1*-containing plasmid of S2568, as this plasmid was not circularized. The dereplicated top hits from each plasmid were collected from PLSDB, and annotated using Prokka version 1.14.6 (Seemann, 2014), as were the *vim-1*-containing plasmids of our isolates. Roary version 1.7.8 (Page et al., 2015) was then used to identify shared genes between all annotated plasmids. The resulting gene presence/absence matrix was used to produce a distance matrix based on Jaccard distances using the philentropy version 0.8.0 (Drost et al., 2018) package in R version 4.3.0 (R Core Team, 2021) which was then used to construct a dendogram using ape version 5.8 (Paradis & Schliep, 2019). The distance matrix was visualized using iTOL (Letunic & Bork, 2021).

### Sequence comparison of OmpC and OmpD porins

All proteins annotated as OmpC or OmpD in the genomes of the six clinical isolates, as well as *E. coli* EcGQ0079 (DH5α with a pBBR1mcs-5::*vim-1* derivative), were extracted from the proteomes. All porins were aligned using the Clustal Omega (Sievers & Higgins, 2017) MBL-EBI webserver (ebi.ac.uk/jdispatcher/msa/clustalo) to generate a multisequence alignment and a percent identity matrix. The untrimmed alignment was used to create a phylogeny using IQ-TREE version 2.2.2.4 (Minh et al., 2020) with the Q.pfam+F+G4 model, as it was identified as the best-scoring model by ModelFinder based on Bayesian information criterion (BIC) and with model search limited to the LG, WAG, JTT, Q.pfam, JTTDCMut, DCMut, VT, PMB, BLOSUM62 and Dayhoff models. Branch supports were assessed using Shimodaira-Hasegawa-like approximate likelihood ratio test (SH-aLRT) (Anisimova & Gascuel, 2006) and an ultrafast bootstrap analysis, with both metrics calculated from 1,000 replicates. The phylogeny and percent identity matrix were then visualized with the iTOL web server (Letunic & Bork, 2021).

### Cloning of *vim-1*

PCR primers MR001 and MR003 (see **Table S1** for all primer sequences) were used to amplify *vim-1* and the preceding *attC* site from *E. coli* F5446 using Q5 DNA polymerase (New England Biolabs [NEB]), which was then purified using a Monarch PCR & DNA Cleanup Kit (NEB). The PCR product and the pBBR1mcs-5 cloning vector (Kovach et al., 1995) were individually digested with both BamHI-HF (NEB) and HindIII-HF (NEB) overnight at 37°C; pBBR1mcs-5 was subsequently dephosphorylated using Quick CIP (NEB). Lastly, the *vim-1*-containing DNA fragments were ligated into pBBR1mcs-5 using T4 DNA ligase (NEB). Ligation products were transformed into chemically competent *E. coli* DH5α and plated on LB with gentamicin and X-gal, and correctly assembled plasmids were preliminarily identified based on blue-white screening. Plasmid DNA was purified from transformants using a Monarch Plasmid DNA Miniprep Kit (NEB) and the *vim-1* sequences then were verified using Sanger sequencing with primers M13-F and M13-R (CHU de Québec-Université Laval Research Center). Plasmids containing *vim-1* were also transformed into electrocompetent cultures of *Klebsiella grimontii* KP5022 (Streicher et al., 1974) and *E. coli* MG1655 (Datsenko & Wanner, 2000) via electroporation and plated on LB with gentamicin to select for the presence of the plasmid.

### Adaptive laboratory evolution (ALE)

Strains of interest were grown overnight in MH broth and diluted to an optical density at 600 nm (OD_600_) of 0.05 in 200 µL MH broth in 96-well microplates. Each microplate contained an ertapenem gradient, where the ertapenem concentration of adjacent wells differed by a factor of two. Microplates were taped closed, inserted into a Synergy H1 plate reader, and incubated for 24 hours at 37°C with shaking. OD_600_ measurements were recorded every 15 minutes. Following 24 hours of incubation, 2 µL from the well with the highest concentration of ertapenem that allowed growth were sub-cultured into each well of an ertapenem concentration gradient in a fresh 96-well plate. This process was repeated until there was a sufficient increase in the concentration of ertapenem tolerated by *vim-1*-containing strains.

To test if the plasmids in the derived isolates were sufficient to produce an ertapenem resistance phenotype, plasmid DNA from the derived isolates was purified with a Monarch Plasmid DNA Miniprep Kit (NEB) and transformed into electrocompetent *E. coli* DH5α cultures via electroporation, then plated on LB with Gm to select for plasmid uptake.

Plasmids from the strains recovered at the end of the ALE were purified using a Monarch Plasmid Miniprep Kit (NEB) and the full plasmid sequence was determined using ONT sequencing by Plasmidsaurus (Louisville, KY, USA). Sequencing results were visualized using the online software Benchling (benchling.com), with alignment performed using MAFFT (Katoh et al., 2019). For one ALE experiment, genomic DNA from an ancestral strain (EcGQ0079) and two derived lineages (EcGQ0088 and EcGQ0089) was purified and sequenced at the Microbial Genome Sequencing Center using an Illumina NextSeq 550 instrument with 150 bp paired-end reads. Illumina reads were then trimmed using BBduk version 38.96 (Bushnell, 2021) and trimmomatic version 0.39 (Bolger et al., 2014). A reference genome was assembled using the Illumina reads of the ancestral strain and Unicycler version 0.5.0 (Wick et al., 2017) with SPAdes version 3.15.4 (Prjibelski et al., 2020) and then annotated using Prokka version 1.14.6 (Seemann, 2014). Next, polymorphisms between the derived isolates and the reference genome were identified using Snippy version 4.6.0 (Seemann, 2015) with bwa version 0.7.17-r1198-dirty (Li & Durbin, 2009). Larger deletions were identified using the samtools version 1.15 coverage and depth functions (Danecek et al., 2021) and the BAM files returned by Snippy.

### Antibiotic susceptibility

Minimum inhibitory concentrations (MIC) of ceftazidime, meropenem, and ertapenem were determined with ThermoFisher Oxoid M.I.C.Evaluator (M.I.C.E.) strips according to the manufacturer’s standard protocol on MH agar plates at 37°C. Antibiotic susceptibility was determined in accordance with CLSI guidelines (CLSI, 2023).

### Reverse transcriptase quantitative PCR (RT-qPCR)

Three biological replicates of each strain of interest were inoculated into 2 mL of Mueller-Hinton broth with relevant antibiotics (10 µg/mL of gentamicin was used for strains containing the pBBR1mcs-5 construct to stabilize the plasmid, and a sub-inhibitory concentration of 0.5 µg/mL of ertapenem was used for clinical isolates to induce *vim-1* expression) and grown overnight at 37°C. The following day, cells were pelleted and washed with fresh media, then sub-cultured to a starting density of OD_600_ = 0.05, grown to a final density of OD_600_ = 0.5 at 37°C, pelleted, flash frozen, and stored at −80°C. RNA was purified from cell pellets with ZymoBIOMICS RNA Miniprep Kit (Zymo Research) according to the manufacturer’s instructions, which was then treated with a TURBO DNA-*free* Kit (Thermo Fisher) following the manufacturer’s instructions, to ensure the samples were free of contaminating DNA. Next, cDNA was synthesized from 2 µg of template RNA using the SuperScript IV VILO Master Mix Kit (Invitrogen), according to the manufacturer’s instructions.

To examine the expression of *vim-1*, qPCR was performed using a BioRad CFX Opus 96 Real-Time PCR System. Expression of *vim-1* was normalized based on expression of the 16S rRNA gene. Standard curves were created for the 16S rRNA gene (primers: SK003 and SK004) and *vim-1* (primers: GD018 and GD019) in 16 µL reactions that included 8 µL of SsoAdvanced Universal SYBR Green qPCR Supermix (BioRad), 187.5 nM of each primer, and between 0.002 and 200 ng of cDNA. The standard curves for both primer sets gave efficiencies >94% and R^2^ values > 0.99 (**Figure S1**). All subsequent qPCRs used 2 ng of cDNA.

### Data availability

The ONT and Illumina data used to generate the whole genome sequences are available through the National Center for Biotechnology Information (NCBI) database under BioProject accession PRJNA1305022. As NCBI had not yet finished processing the annotated genomes by the time this manuscript was uploaded to NCBI, copies of the annotated genome files have been uploaded to GitHub (github.com/diCenzo-Lab/015_2025_VIM-1_analyses), and will be made available via NCBI under BioProject accession PRJNA1305022 once processing is completed. All code to repeat the computational analyses reported in this study is available through GitHub (github.com/diCenzo-Lab/015_2025_VIM-1_analyses).

## RESULTS

### Whole genome sequencing of six *vim-1*-positive *Enterobacterales* clinical isolates

Six multidrug resistant clinical isolates of order *Enterobacterales* were isolated from patients at the Kingston Health Sciences Center, Kingston, Ontario, Canada between 2015 and 2020. Despite all six isolates testing positive for the presence of *vim-1* based on a PCR screen, they displayed a wide range of resistance to the carbapenem antibiotics ertapenem and meropenem, with some isolates phenotypically testing as susceptible despite the presence of *vim-1* (**Table 2**). In contrast, all strains were equally and highly resistant to the cephalosporin antibiotic ceftazidime (**Table 2**).

**Table 2.** Minimum inhibitory concentrations of ceftazidime, meropenem, and ertapenem against six clinical isolates, together with the copy numbers of *vim-1*-containing plasmids and *vim-1* expression levels.

| Isolate | Minimum inhibitory concentration (MICs; µg/mL) * |  |  | *vim-I* plasmid<br>copy number † | *vim-I* expression<br>(ΔCq) ¥ |
| --- | --- | --- | --- | --- | --- |
|  | Ceftazidime | Meropenem | Ertapenem |  |  |
| *E. coli* F5446 | ≥256 (R) | 6 (R) | 2 (R) | 1 | 11.8 ± 1.0 |
| *E. coli* S2568 | ≥256 (R) | 0.75 (S) | 3 (R) | 15 | 13.4 ± 0.6 |
| *E. hormaechei* F48994 | ≥256 (R) | 4 (R) | 2 (R) | 2 | 10.5 ± 0.1 |
| *E. hormaechei* H17629 | ≥256 (R) | 0.38 (S) | 0.25 (S) | 1 | 13.4 ± 0.6 |
| *E. hormaechei* H70375 | ≥256 (R) | 0.5 (S) | 0.38 (S) | 5 | 9.6 ± 0.7 |
| *K. pneumoniae* T64870 | ≥256 (R) | 1 (I) | 0.25 (S) | 4 | 11.1 ± 0.4 |

\* MICs are marked (S) susceptible, (I) intermediate, or (R) resistant according to the CLSI guidelines.
† Copy number of the *vim-I*-containing plasmids are based on the copy numbers reported by the Flye assembler.
¥ Values represent the average ΔCq of three biological replicates ± standard deviation; lower numbers represent higher expression. Expression of *vim-I* was normalized based on expression of the 16S rRNA genes.

Whole genome sequencing, assembly, and annotation was performed for all six clinical isolates. The resulting assemblies were high quality (completeness >99% and contamination <1% as determined by CheckM) and consisted of between three and seven contigs (**Table 3**). For all six strains, the chromosome was assembled into a single, circular replicon. In addition, three of the six genomes appeared to be complete, closed genomes, while the other three included at least one plasmid split into two contigs. Taxonomic classification of the isolates indicated that F48994, H17629, and H70375 belonged to the species *Enterobacter hormaechei,* F5446 and S2568 belonged to the species *Escherichia coli,* and T64870 belonged to the species *Klebsiella pneumoniae*.

**Table 3.** Genome assembly and annotation statistics.

| Characteristic | *E. coli*<br>F5446 | *E. coli*<br>S2568 | *E. hormaechei*<br>F48994 | *E. hormaechei*<br>H17629 | *E. hormaechei*<br>H70375 | *K. pneumoniae*<br>T64870 |
| --- | --- | --- | --- | --- | --- | --- |
| Genome size (bp) | 5,382,174 | 5,388,229 | 5,160,253 | 4,988,147 | 5,240,385 | 5,768,527 |
| No. of scaffolds | 4 | 5 | 7 | 3 | 6 | 3 |
| G+C content (%) | 50.73% | 50.75% | 54.77% | 55.18% | 54.49% | 57.02% |
| No. of protein coding genes | 4,878 | 4,884 | 4,731 | 4,450 | 4,798 | 5,327 |
| Scaffolds N50 | 1 | 1 | 1 | 1 | 1 | 1 |
| Scaffolds L50 | 5,028,008 | 5,044,934 | 4,722,177 | 4,794,996 | 4,680,518 | 5,452,037 |
| Completeness (%) | 99.97 | 99.97 | 99.87 | 99.71 | 99.51 | 99.84 |
| Contamination (%) | 0.25 | 0.06 | 0.22 | 0.98 | 0.82 | 0.33 |

Each of the isolates contained at least six perfect hits (100% nucleotide sequence identity) to different ARGs (**Table S2**). As expected, all six isolates carried the *vim-1* gene, but they also contained between 0 and 6 additional β-lactamase genes of the types ACT-24, CTX-M-15, LAP-2, OXA-1, OXA-9, TEM-1, SHV-12, and SHV-28 (**Table S2**). The β-lactamases ACT-24, CTX-M-15, LAP-2, and TEM-1 are not known to interact with carbapenems, and therefore should not contribute to ertapenem or meropenem resistance (Alcock et al., 2020). Likewise, while some OXA-family β-lactamases are considered ESBLs, OXA-1 and OXA-9 are not known to interact with carbapenems (Evans & Amyes, 2014), and SHV-12 (found in isolates F5446 and S2568) and SHV-28 (found in isolate T64870) are only capable of mediating carbapenem resistance in combination with major outer membrane porin modifications (Leavitt et al., 2009). Although these additional β-lactamase genes likely contributed to high ceftazidime resistance in the clinical isolates, these results suggest that *vim-1* is primarily responsible for the observed carbapenem resistance.

### Genomic context of the *vim-1* genes of the six clinical isolates

As expected, *vim-1* was situated within an integron in each of the six clinical isolates. However, the type of integron on which *vim-1* was housed differed along species lines; the *E. coli* isolates contain In916 integrons while the other isolates contain In110 integrons (Matsumura et al., 2017). In916 and In110 are clinical class 1 integrons (**Figure 1**) that both contain a 5’ conserved sequence that includes the *intI1* integrase, a 3’ conserved sequence that includes the ARGs *qacEΔ1* and *sul1*, and a variable region encoding additional ARGs and that differs in content between In916 and In110 (Alcock et al., 2020). Notably, in all six of our strains, *vim-1* is the first gene within the variable region.

**Figure 1.**
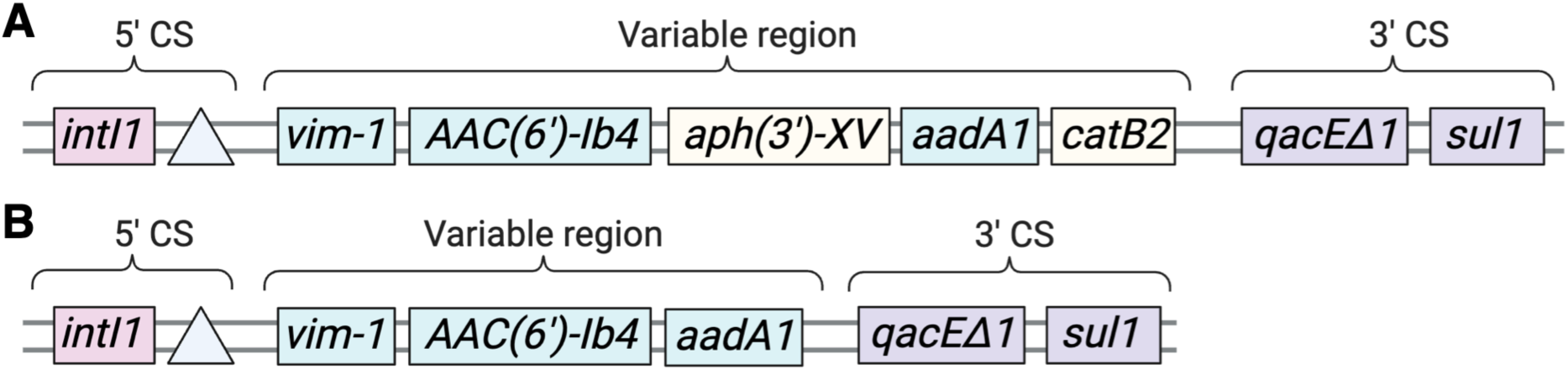
The structure of the *vim-1*-containing class 1 integrons identified in this study. **(A)** *Escherichia coli* F5446 and *E. coli* S2568 encode *vim-1* within In916 integrons, whereas **(B)** *Enterobacter hormaechei* F48994, *E. hormaechei* H17629, *E. hormaechei* H70375, and *Klebsiella pneumoniae* T64870 encode *vim-1* within In110 integrons. (**A**, **B**) The 5’ conserved sequences (5’ CS), variable regions, and 3’ conserved sequences (3’ CS) are indicated above each diagram. The position of *attL1* is represented as a light blue triangle, while the two genes present in In916 but absent from In110 are shown in light yellow. The diagrams are not drawn to scale.

All six *vim-*1-containing integrons were housed on plasmids rather than the bacterial chromosome. *E. coli* clinical isolates F5446 and S2568 contained *vim-1* in the context of IncA plasmids, while other isolates contained their *vim-1* in the context of IncN (*E. hormaechei* H17629), IncR (*E. hormaechei* H70375), or multi-locus IncN-IncR plasmids (*E. hormaechei* F48994, *K. pneumoniae* T64870). The copy number (as determined by the genome assembler Flye) of *vim-1-*containing plasmids varied across clinical isolates, but these variations did not positively correlate with increases in *vim-1* expression or tolerance to carbapenem antibiotics (**Table 2**). Likewise, relative *vim-1* expression, as determined by RT-qPCR, was not positively correlated with relative carbapenem resistance (**Table 2**).

### Relationships between the *vim-*1-containing plasmids of the six clinical isolates

To explore the introduction and dissemination of *vim-1* genes within Kingston (Ontario, Canada) and the surrounding region, a mid-point rooted, bifurcating dendrogram based on shared gene content was produced for the *vim-1* containing plasmids of our six clinical isolates together with a set of 75 related plasmids (**Figure 2**). Of the 75 external plasmids that were included in the analysis, 24 carried *vim-1*. Overall, the *vim-1* plasmids from our clinical isolates formed three distinct clades that also included *vim-1*-containing plasmids from isolates collected in Toronto (Ontario, Canada), suggesting three separate introductions of *vim-1* genes into the Kingston region and the spread of these genes/pathogens between Toronto and Kingston.

**Figure 2.**
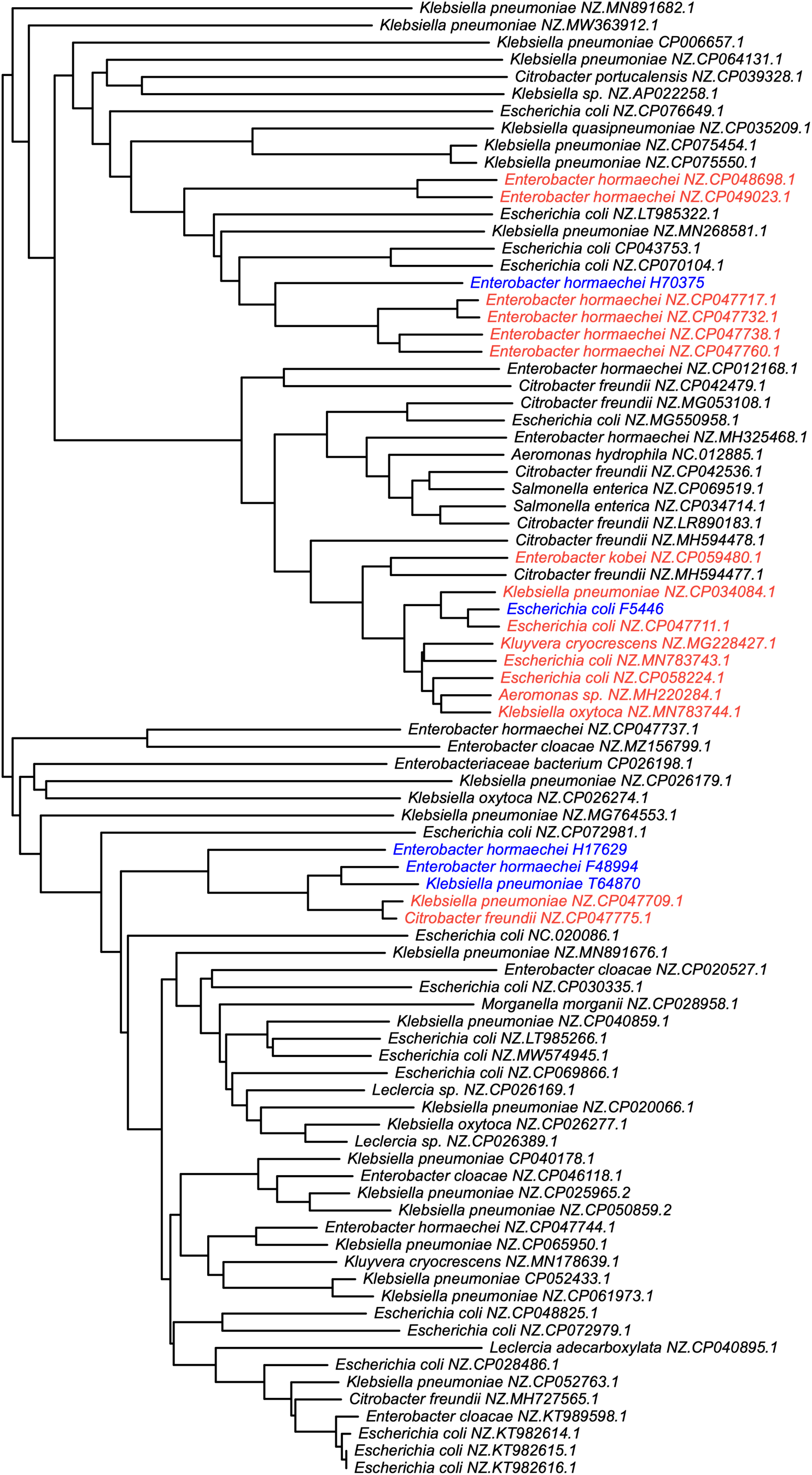
Relationships between the *vim-1*-containing plasmids of this study and related plasmids of the plasmid database (PLSDB). Plasmids related to the *vim-1*-containing plasmids identified in the six focal clinical isolates of this study were identified in PLSDB using blast, and a dendogram was constructed from a distance matrix built on gene presence/absence data. Shown in blue are the *vim-1*-containing plasmids identified in the current study, while red is used to represent other *vim-1* containing plasmids. Plasmids are named according to the species in which the plasmid was identified, followed by the GenBank accession of the plasmid.

The *vim-1*-containing plasmids of *E. hormaechei* F48994, *E. hormaechei* H17629, and *K. pneumoniae* T64870 (which all contain *vim-1* within an In110 integron) formed a clade with two other *vim-1*-containing plasmids: pKpn13-2 (CP047709) and pCfr13-2 (CP047775). These plasmids were isolated from rectal samples of *K. pneumoniae* and *Citrobacter freundii,* respectively, by the Toronto Invasive Bacterial Diseases Network (TIBDN) from a single patient in Toronto, ON (Kohler et al., 2020). These results suggest that this *vim-1*-containing plasmid has spread between pathogens through horizontal transfer within southern Ontario. Notably, pKpn13-2 and pCfr13-2 are nested within the F48994, H17629, and T64870 cluster, suggesting that the detection of this plasmid in Toronto may have been due to spread from Kingston.

Four *vim-1-*containing plasmids clustered with the *E. hormaechei* H70375 plasmid (which contains *vim-1* within an In110 integron): pEclE2-2 (CP047717), pEcl6-3 (CP047732), pEcl5-2 (CP047738), and pEcl2-3 (CP047760). These plasmids all came from *E. hormaechei* isolates, one of which (pEclE2-2) was an environmental isolate collected from Toronto, ON sewage water in 2015, while the others are clinical isolates collected from hospitals by the TIBDN in Toronto, ON between 2011 and 2014 (Kohler et al., 2020). These plasmids all have IncR replicons and are thought to share a common ancestral IncR plasmid (Kohler et al., 2020). The plasmid from H70375 being the deepest branching lineage within this cluster, suggests that the detection of these *vim-1*-containing plasmids in Toronto may have been the result of its spread from Kingston.

Finally, there is a large cluster of *vim-1*-containing plasmids that includes the *vim-1*-containing plasmids of *E. coli* F5446 and S2568 (which contain *vim-1* within an In916 integrons); the plasmid of S2568 is not shown in the dendrogram due to it being nearly identical to that of F5446 but not fully assembled. The most closely related plasmid to that from *E. coli* F5446 is pEco15-1 (CP047711), which was isolated from *E. coli* from a rectal swab gathered by the TIBDN in 2015 (Kohler et al., 2020). Interestingly, the other plasmids within this cluster are not local to Ontario and come from isolates across the globe and from multiple species. These include pR210-2 (CP034084) from a *K. pneumoniae* isolate collected at the Hong Kong Polytechnic University (unpublished), pRIVM0001_VIM-1_171012_B12 (MH220284) from a Dutch *Aeromonas* sp. sample (unpublished), p550_IncA_VIM_1 (CP058224) from an Italian *E. coli* sample (Mattioni Marchetti et al., 2020), pGA_VIM (MN783743) from an Italian *E. coli* sample (Arcari et al., 2020), pKC-BO-N1-VIM (MG228427) from an Italian *Kluyvera cryocrescens* sample (Gaibani et al., 2018), and pFDL_VIM (MN783744) from an Italian *Klebsiella oxytoca* sample (Arcari et al., 2020).

### Expression of *vim-1* alone facilitates limited resistance to ertapenem in laboratory strains

To explore the contribution of *vim-1* to the carbapenem antibiotic ertapenem, *vim-1* of *E. coli* F5446 was PCR amplified and expressed under the control of the pLac promoter on the plasmid pBBR1mcs-5 in three non-pathogenic lab strains: *E. coli* DH5α, *E. coli* MG1655, and *K. grimontii* KP5022. Antibiotic susceptibility tests of *E. coli* DH5α, *E. coli* MG1655, and *K. grimontii* KP5022 strains carrying an empty pBBR1mcs-5 vector confirmed that the vector itself did not contribute to ertapenem, meropenem, or ceftazidime resistance (**Table 4**). On the other hand, introduction of pBBR1mcs-5::*vim-1* resulted in an ∼ 20-fold to 64-fold increase in resistance to the cephalosporin antibiotic ceftazidime (**Table 4**), confirming *vim-1* was expressed and functional in all strains. Introduction of pBBR1mcs-5::*vim-1* into *K. grimontii* KP5022 also resulted in an ∼ 40-fold increase in resistance to the carbapenem meropenem but had little impact on resistance to the carbapenem ertapenem (**Table 4**). Similarly, introduction of *vim-1* into the two *E. coli* strains resulted in at most a 3-fold increase in resistance to meropenem and ertapenem (**Table 4**). In all cases, introduction of *vim-1* did not result in any of the three strains being classified as resistant to any of the three antibiotics according to the CLSI guidelines.

**Table 4.** Minimum inhibitory concentrations of ceftazidime, meropenem, and ertapenem against laboratory strains expressing *vim-1*, both before and after repeated passaging in growth media containing ertapenem.

| Species | Strain | Description | Minimum inhibitory concentration (MICs; µg/mL) * |  |  |
| --- | --- | --- | --- | --- | --- |
|  |  |  | Ceftazidime | Meropenem | Ertapenem |
| *E. coli* | DH5α | Wild type | 0.094 (S) | 0.016 (S) | 0.006 (S) |
| *E. coli* | J243 | DH5α carrying pBBR1mcs-5 | 0.094 (S) | 0.016 (S) | 0.004 (S) |
| *E. coli* | EcGQ0033 | DH5α carrying pBBR1mcs-5::*vim-1* | 2 (S) | 0.032 (S) | 0.006 (S) |
| *E. coli* | MG1655 | Wild type | 0.19 (S) | 0.023 (S) | 0.012 (S) |
| *E. coli* | EcGQ0110 | MG1655 carrying pBBR1mcs-5 | 0.19 (S) | 0.032 (S) | 0.012 (S) |
| *E. coli* | EcGQ0036 | MG1655 carrying pBBR1mcs-5::*vim-1* | 8 (I) | 0.094 (S) | 0.023 (S) |
| *K. grimontii* | KP5022 | Wild type | 0.125 (S) | 0.023 (S) | 0.012 (S) |
| *K. grimontii* | KpGQ0020 | KP5022 carrying pBBR1mcs-5 | 0.125 (S) | 0.032 (S) | 0.006 (S) |
| *K. grimontii* | KpGQ0001 | KP5022 carrying pBBR1mcs-5::*vim-1* | 8 (I) | 0.94 (S) | 0.012 (S) |
| *K. grimontii* | KpGQ0008 | KpGQ0001 ALE derived isolate | >256 (R) | 1.5 (I) | 0.004 (S) |
| *K. grimontii* | KpGQ0016 | DH5α with pBBR1mcs-5::*vim-1* of KpGQ0008 | >256 (R) | 1.0 (S) | 0.006 (S) |
| *K. grimontii* | KpGQ0009 | KpGQ0001 ALE derived isolate | >256 (R) | 1.5 (I) | 0.006 (S) |
| *K. grimontii* | KpGQ0017 | DH5α with pBBR1mcs-5::*vim-1* of KpGQ0009 | >256 (R) | 1.0 (S) | 0.006 (S) |
| *E. coli* | EcGQ0082 | EcGQ0033 ALE derived isolate | >256 (R) | 0.25 (S) | 0.006 (S) |
| *E. coli* | EcGQ0141 | DH5α with pBBR1mcs-5::*vim-1* of EcGQ0082 | >256 (R) | 0.25 (S) | 0.064 (S) |
| *E. coli* | EcGQ0137 | EcGQ0082 cured of pBBR1mcs-5::*vim-1* | 0.5 (S) | 0.047 (S) | 0.004 (S) |
| *E. coli* | EcGQ0083 | EcGQ0033 ALE derived isolate | >256 (R) | 0.23 (S) | 0.032 (S) |
| *E. coli* | EcGQ0142 | DH5α with pBBR1mcs-5::*vim-1* of EcGQ0083 | >256 (R) | 0.25 (S) | 0.047 (S) |
| *E. coli* | EcGQ0138 | EcGQ0083 cured of pBBR1mcs-5::*vim-1* | 1 (S) | 0.047 (S) | 0.012 (S) |
| *E. coli* | EcGQ0079 | DH5α with pBBR1mcs-5::*vim-1* of KpGQ0009 | >256 (R) | 0.094 (S) | 0.016 (S) |
| *E. coli* | EcGQ0088 | EcGQ0079 ALE derived isolate | >256 (R) | >32 (R) | >32 (R) |
| *E. coli* | EcGQ0089 | EcGQ0079 ALE derived isolate | >256 (R) | >32 (R) | >32 (R) |
| *E. coli* | EcGQ0143 | DH5α with pBBR1mcs-5::*vim-1* of EcGQ0088 | >256 (R) | 0.19 (S) | 0.064 (S) |
| *E. coli* | EcGQ0144 | DH5α with pBBR1mcs-5::*vim-1* of EcGQ0089 | >256 (R) | 0.25 (S) | 0.064 (S) |
| *E. coli* | EcGQ0139 | EcGQ0088 cured of pBBR1mcs-5::*vim-1* | 0.25 (S) | 0.094 (S) | 0.25 (S) |
| *E. coli* | EcGQ0140 | EcGQ0089 cured of pBBR1mcs-5::*vim-1* | ND | ND | ND |

ND – Not determined due to poor growth of the strain.
\* MICs are marked (S) susceptible, (I) intermediate, or (R) resistant according to the CLSI guidelines.

To increase the level of resistance to ertapenem, adaptive laboratory evolution (ALE) experiments were performed with *K. grimontii* KP5022 (pBBR1mcs-5::*vim-1*) and *E. coli* DH5α (pBBR1mcs-5::*vim-1*). Replicates of each strain were grown in the presence of increasingly higher concentrations of ertapenem over a span of five days, following which individual isolates were isolated from two of the replicates of each species. Antibiotic susceptibility testing indicated that all derived lineages had extremely high resistance to ceftazidime but little to no increase in resistance to meropenem or ertapenem under the tested conditions (**Table 4**).

Transfer of the pBBR1mcs-5::*vim-1* plasmids from the *K. grimontii* or *E. coli* ALE derived isolates to otherwise wildtype *K. grimontii* KP5022 or *E. coli* DH5α, respectively, demonstrated that the mutation(s) responsible for the elevated ceftazidime resistance in the derived isolates are linked to the plasmids (**Table 4**). To identify the responsible mutation(s), we used ONT sequencing to determine the complete sequence of the pBBR1mcs-5::*vim-1* plasmids from the derived isolates and compared them to the sequence of the original pBBR1mcs-5::*vim-1* plasmid. Unexpectedly, we observed that the plasmids isolated from the derived lineages were all approximately twice the length of the original pBBR1mcs-5::*vim-1* construct, with all genes (including *vim-1*) duplicated. This result suggested that the increased ceftazidime resistance of the derived lineages was a result an increase in *vim-1* gene dosage and thus higher *vim-1* expression. Indeed, RT-qPCR indicated that *vim-1* expression in the derived isolates was ∼ 3.7- to 6.3-fold higher than in the ancestral strains (**Table 5**).

**Table 5.** Expression of *vim-1* in laboratory strains expressing *vim-1*, both before and after repeated passaging in growth media containing ertapenem.

| Species | Strain | Description | *vim-I* expression ($\Delta\Delta Cq$)* |
| --- | --- | --- | --- |
| *K. grimontii* | KpGQ0001 | KP5022 carrying pBBR1mcs-5::*vim-I* | $0.0 \pm 1.0$ |
| *K. grimontii* | KpGQ0008 | KpGQ0001 ALE derived isolate | $2.4 \pm 0.2$ |
| *K. grimontii* | KpGQ0009 | KpGQ0001 ALE derived isolate | $2.6 \pm 0.4$ |
| *E. coli* | EcGQ0033 | DH5α carrying pBBR1mcs-5::*vim-I* | $0.0 \pm 0.3$ |
| *E. coli* | EcGQ0082 | EcGQ0033 ALE derived isolate | $1.9 \pm 0.5$ |
| *E. coli* | EcGQ0083 | EcGQ0033 ALE derived isolate | $2.3 \pm 0.3$ |

\* Values represent the average $\Delta\Delta Cq$ of three biological replicates ± standard deviation. Expression of *vim-I* was normalized based on expression of the 16S rRNA genes and then compared to the average $\Delta Cq$ of KpGQ0001 (for the *K. grimontii* strains) or EcGQ0036 (for the *E. coli* strains). The $\Delta\Delta Cq$ values represent the log(2) fold change compared to the reference. The difference in expression of *vim-I* in each ALE was statistically significant based on t-tests (p-value < 0.05).

### *E. coli* DH5α resistance to ertapenem requires both *vim-1* and mutations of an outer membrane porin

Despite the increased *vim-1* expression in the evolved lineages described above, none of the derived isolates were considered resistant to either ertapenem or meropenem under the tested conditions based on the CLSI guidelines (**Table 4**). To evolve higher ertapenem resistance in *E. coli* DH5α, an *E. coli* DH5α strain carrying the enlarged pBBR1mcs-5::*vim-1* plasmid isolated from one of the *K. grimontii* KP5022 derived lineages was subjected to a six-day ALE in the presence of ertapenem, following which individual isolates were collected from two replicate cultures. Antibiotic susceptibility testing confirmed that the derived isolates displayed very high resistance to all three tested antibiotics (ceftazidime, meropenem, and ertapenem) (**Table 4**).

Curing of the *vim-1*-containing plasmid from the derived lineages resulted in close to ancestral levels of resistance, while transfer of the *vim-1*-containing plasmid to otherwise wildtype *E. coli* DH5α resulted in little resistance to meropenem or ertapenem (**Table 4**). Together, these results suggest that the high ertapenem resistance of the derived lineages is dependent both on the presence of *vim-1* and at least one additional mutation on the chromosome. To identify the chromosomal mutations responsible for elevated ertapenem resistance, whole genome sequencing of two derived isolates (strains EcGQ0088 and EcGQ0089; **Table 4**), as well as the ancestral strain (strain EcGQ0079; **Table 4**), was performed. Mapping of the EcGQ0088 sequencing reads to the EcGQ0079 reference genome identified a single polymorphism: an in-frame deletion of six nucleotides within *ompC*, which encodes outer membrane porin C, a transmembrane protein that facilitates the transport of beta-lactam antibiotics across the outer membrane (Jaffe et al., 1982). Similarly, EcGQ0089 contained only a single mutation: a 26 kb deletion that spanned *ompC*, among other genes. Overall, these results indicate that high levels of *vim-1*-mediated ertapenem resistance in *E. coli* DH5α is dependent on loss-of-function mutations within *ompC*.

## DISCUSSION

### The gene *vim-1* is disseminated clonally and by plasmid-mediated HGT in Ontario

Plasmids from all clinical isolates identified here can be linked to those associated with hospital outbreaks in Ontario and Europe and appear to form three distinct clades. The first clade, which forms around the *vim-1* plasmids of *E. coli* F5664 and S2568, contains plasmids belonging to the IncA incompatibility group. The spread of plasmids in this clade, both geographically and across species of bacteria, suggests plasmids of this cluster are globally distributed and have been spread through HGT. The second clade contained the *vim-1*-containing plasmids of *E. hormaechei* H17629, F48994, *K. pneumoniae* T64870, and two additional plasmids belonging to the multilocus IncR-IncN incompatibility group from Toronto (Kohler et al., 2020). The third clade was formed around the *vim-1-*containing plasmid from *E. hormaechei* H70375 and four additional plasmids from Toronto (Kohler et al., 2020), all belonging to the IncR incompatibility group. For both of these clades, the detection of similar plasmids in isolates from two cities within Ontario, Canada, but not yet elsewhere at the time this analysis was performed, suggests that these plasmids have been spread across species via HGT within Ontario. However, it is unknown whether IncR plasmids are transferable. Their broad host range suggests that they should be transferable (Bielak et al., 2011; Kohler et al., 2020), but this plasmid complex lacks a transfer system and relaxase necessary for mobilization (Compain et al., 2014; Kohler et al., 2020; Smillie et al., 2010). Therefore, it is possible that plasmids in the third clade can be found in *E. hormaechei* isolates across Ontario due to vertical transfer and subsequent evolution, but such conclusions would require further phylogenomic analysis. Overall, these findings support the current consensus on *vim-1* in Ontario; that the gene is spread both clonally and by plasmid-mediated HGT (Jamal et al., 2021; Kohler et al., 2020; Tijet et al., 2013).

### Genomic background influences *vim-1* activity

Among the six clinical isolates, the MICs of ertapenem ranged from 0.25 μg/mL, which is considered sensitive by CLSI guidelines, to 3 μg/mL, which is considered resistant. Ertapenem resistance was even lower (0.006 to 0.012 µg/mL) when *vim-1* was cloned and introduced into three common *Enterobacteriaceae* laboratory strains. This phenotypic variation is consistent with previous studies noting phenotypic variability of MBLs when introduced into diverse bacterial hosts (Socha et al., 2019). The differences in resistance phenotype do not fall cleanly around species lines, since there are variations in MICs of ertapenem and meropenem between individual *E. coli* and *E. hormaechei* isolates, although in general, the *E. coli* clinical isolates show higher ertapenem resistance than the other four isolates.

The above observations led us to consider possible mechanistic explanations for the phenotypic variability across our six clinical isolates. Since *vim-1* is integron-encoded and appears in identical positions relative to the integron promoter in all six clinical isolates, differences in promoter proximity and promoter strength are unlikely to explain the differences in resistance phenotypes conferred by *vim-1*; indeed, no obvious relationship was observed between resistance and *vim-1* expression as measured by RT-qPCR. However, mRNA abundance and protein periplasmic concentrations are not necessarily correlated, and it has been noted that variation in MBL periplasmic concentration contributes to the phenotypic variation of MBLs across strains (Socha et al., 2019).

Another possibility is that the variations in VIM-1-mediated ertapenem resistance across the strains is due to epistatic interactions between *vim-1* and other genomic loci. Previous studies have observed that high resistance to ertapenem and other carbapenems depends on the presence of a β-lactamase paired with reduced outer membrane permeability (Codjoe et al., 2025; Doumith et al., 2009; Jacoby et al., 2004; Leavitt et al., 2009; Szabó et al., 2006; Woodford et al., 2007). With one exception (Codjoe et al., 2025), those studies focused on variations in membrane permeability in strains with ESBLs rather than strains with true carbapenemases (like VIM-1). Examining the genomes of our six clinical isolates for outer membrane porins revealed that four of the six strains had two copies of *ompC*, and that the *E. hormaechei* strains also carried *ompD*. In addition, the OmpC amino acid sequences of our six isolates were highly variable, sharing between 62.9% and 100% sequence identity (**Figure S2**). Thus, we hypothesize that variation in membrane permeability contributed to the differences in ertapenem resistance observed in our clinical isolates, which would suggest that reduced membrane permeability is likely an important factor impacting carbapenem resistance even in isolates with true carbapenemases.

### VIM-1 alone was not sufficient to confer carbapenem resistance

To experimentally explore factors influencing VIM-1-mediated ertapenem resistance, *vim-1* was amplified from a *vim-1*^+^ clinical isolate without its native promoter and expressed on a plasmid in the laboratory strains *E. coli* DH5α, MG1655, and *K. grimontii* KP5022. For all strains, the presence of *vim-1* alone was insufficient to confer resistance to the carbapenem antibiotics meropenem and ertapenem, but did confer resistance to the cephalosporin antibiotic ceftazidime. These results confirmed *vim-1* was expressed and the encoded enzyme was active and suggested that *vim-1* alone is a strong resistance determinant for cephalosporin antibiotics (like ceftazidime), but not necessarily for carbapenems like ertapenem and meropenem.

ALE was used to identify mutations increasing ertapenem resistance in the laboratory strains carrying *vim-1*. In all cases, the primary mutation that we identified was an increase in the *vim-1* copy number, and thus expression. Although the increase in *vim-1* expression increased the concentration of ertapenem that the strains could survive in the liquid-based ALE experiments, this mutation had little impact on ertapenem MIC as determined by plate-based assays. We hypothesize that the difference between the liquid-based and agar-based assays reflects that in the liquid-assays, the concentration of ertapenem constantly decreased as VIM-1 degraded the antibiotic, allowing growth when the concentration was sufficiently low. On the other hand, the concentration of ertapenem may not have changed significantly in the plate-based assay due to diffusion of the antibiotic throughout the plate.

On the other hand, strong VIM-1-mediated ertapenem resistance in the plate-based assay was observed when *ompC*, encoding an outer membrane porin, was mutated or deleted, although these mutations came with a trade-off of reduced growth in the absence of antibiotics. This is consistent with previous studies showing synergistic effects of pairing an ESBL with reduced outer membrane permeability (Codjoe et al., 2025; Doumith et al., 2009; Jacoby et al., 2004; Szabó et al., 2006; Woodford et al., 2007). Porins are pore-forming proteins with a β-barrel structure that allow for the passive transport of hydrophilic compounds across the bacterial outer membrane (Schulz, 2002). OmpC is a non-specific porin that plays a role in both membrane integrity and antibiotic transport (Choi & Lee, 2019), meaning its mutation or deletion would result in reduced transport of β-lactam antibiotics across the outer membranes, and reduce their concentration in the periplasm to a level that does not overwhelm VIM-1.

### Conclusions

Here, we describe six *Enterobacterales* clinical isolates carrying VIM-1 on horizontally transmissible plasmids. These plasmids appear to be globally distributed and to have spread both clonally and via plasmid-mediated HGT. In addition, the six isolates vary in carbapenem resistance, which we hypothesize is driven primarily by variations in outer membrane permeability. In support of this, we observed that transfer of *vim-1* to common laboratory strains failed to result in ertapenem resistance unless paired with mutation or deletion of *ompC* encoding an outer membrane porin. This result is consistent with studies of strains expressing ESBLs, although the requirement for reduced outer membrane permeability for VIM-1-mediated ertapenem resistance has been less studied. Finally, these results further highlight that PCR-based detection of *vim-1* is not necessarily sufficient to demonstrate that an isolate is resistant to carbapenems without functional verification.

## ACKNOWLEDGEMENTS

This work was supported by the Natural Sciences and Engineering Research Council of Canada (NSERC) through a Discovery Grant to G.C.D. (RGPIN-2020-0700).

## REFERENCES

Alcock, B. P., Raphenya, A. R., Lau, T. T. Y., Tsang, K. K., Bouchard, M., Edalatmand, A., Huynh, W., Nguyen, A.-L. V., Cheng, A. A., Liu, S., Min, S. Y., Miroshnichenko, A., Tran, H.-K., Werfalli, R. E., Nasir, J. A., Oloni, M., Speicher, D. J., Florescu, A., Singh, B., Faltyn, M., Hernandez-Koutoucheva, A., Sharma A. N., Bordeleau, E., Pawlowski, A. C., Zubyk, H. L., Dooley, D., Griffiths, E., Maguire, F., Winsor, G. L., Beiko, R. G., Brinkman, F. S. L., Hsiao, W. W. L., Domselaar, G. V. & McArthur, A. G. (2020). CARD 2020: Antibiotic resistome surveillance with the comprehensive antibiotic resistance database. Nucleic Acids Res, 48(D1), D517–D525. 10.1093/nar/gkz935

Anisimova, M., & Gascuel, O. (2006). Approximate likelihood-ratio test for branches: A fast, accurate, and powerful alternative. Syst Biol, 55(4), 539–552. 10.1080/10635150600755453

Arcari, G., Di Lella, F. M., Bibbolino, G., Mengoni, F., Beccaccioli, M., Antonelli, G., Faino, L., & Carattoli, A. (2020). A multispecies cluster of VIM-1 carbapenemase-producing *Enterobacterales* linked by a novel, highly conjugative, and broad-host-range IncA plasmid forebodes the reemergence of VIM-1. Antimicrob Agents Chemother, 64(4), e02435–19. 10.1128/AAC.02435-19

Bielak, E., Bergenholtz, R. D., Jørgensen, M. S., Sørensen, S. J., Hansen, L. H., & Hasman, H. (2011). Investigation of diversity of plasmids carrying the *bla*TEM-52 gene. J Antimicrob Chemother, 66(11), 2465–2474. 10.1093/jac/dkr331

Bolger, A. M., Lohse, M., & Usadel, B. (2014). Trimmomatic: a flexible trimmer for Illumina sequence data. Bioinformatics, 30(15), 2114–2120. 10.1093/bioinformatics/btu170

Botelho, J., Grosso, F., Quinteira, S., Brilhante, M., Ramos, H., & Peixe, L. (2018). Two decades of *bla*VIM-2-producing *Pseudomonas aeruginosa* dissemination: An interplay between mobile genetic elements and successful clones. Journal Antimicrob Chemother, 73(4), 873–882. 10.1093/jac/dkx517

Bushnell B. (2021). BBMap: BBMap short read aligner, and other bioinformatic tools. [Computer software] https://sourceforge.net/projects/bbmap/

Camacho, C., Coulouris, G., Avagyan, V., Ma, N., Papadopoulos, J., Bealer, K., & Madden, T.L. (2009). BLAST+: architecture and applications. BMC Bioinform, 10(421). 10.1186/1471-2105-10-421

Carattoli, A., Zankari, E., García-Fernández, A., Voldby Larsen, M., Lund, O., Villa, L., Møller Aarestrup, F., & Hasman, H. (2014). In silico detection and typing of plasmids using PlasmidFinder and plasmid multilocus sequence typing. Antimicrob Agents Chemother, 58(7), 3895–3903. 10.1128/AAC.02412-14

Choi, U., & Lee, C.-R. (2019). Distinct roles of outer membrane porins in antibiotic resistance and membrane integrity in Escherichia coli. Front Microbiol, 10(2019). https://www.frontiersin.org/articles/10.3389/fmicb.2019.00953

CLSI. (2023). CLSI M100-ED33:2023 Performance Standards for Antimicrobial Susceptibility Testing (33rd Edition).

Codjoe, F. S., Kotey, F. C. N., & Donkor, E. S. 2025. Profile of outer membrane proteins of carbapenem-resistant Gram-negative bacilli in Ghana. BMC Res Notes, 18(49). 10.1186/s13104-024-07070-6

Compain, F., Frangeul, L., Drieux, L., Verdet, C., Brisse, S., Arlet, G., & Decré, D. (2014). Complete nucleotide sequence of two multidrug-resistant IncR plasmids from *Klebsiella pneumoniae*. Antimicrob Agents Chemother, 58(7), 4207–4210. 10.1128/AAC.02773-13

Council of Canadian Academies. (2019). When Antibiotics Fail: The Expert Panel on the Potential Socio-Economic Impacts of Antimicrobial Resistance in Canada. http://www.deslibris.ca/ID/10102747

Cowie, A., Cheng, J., Sibley, C. D., Fong, Y., Zaheer, R., Patten, C. L., Morton, R. M., Golding, G. B., & Finan, T. M. (2006). An integrated approach to functional genomics: construction of a novel reporter gene fusion library for *Sinorhizobium meliloti*. Appl Environ Microbiol, 72(11), 7156–7167. 10.1128/AEM.01397-06

Currie, C. J., Berni, E., Jenkins-Jones, S., Poole, C. D., Ouwens, M., Driessen, S., Voogd, H. de, Butler, C. C., & Morgan, C. L. (2014). Antibiotic treatment failure in four common infections in UK primary care 1991-2012: Longitudinal analysis. BMJ, 349, g5493. 10.1136/bmj.g5493

Dadgostar, P. (2019). Antimicrobial resistance: Implications and costs. Infect Drug Resist, 12, 3903–3910. 10.2147/IDR.S234610

Danecek, P., Bonfield, J. K., Liddle, J., Marshall, J., Ohan, V., Pollard, M. O., Whitwham, A., Keane, T., McCarthy, S. A., Davies, R. M., & Li, H. (2021). Twelve years of SAMtools and BCFtools. GigaScience, 10(2), giab008. 10.1093/gigascience/giab008

Darling, A. C. E., Mau, B., Blattner, F. R., & Perna, N. T. (2004). Mauve: Multiple alignment of conserved genomic sequence with rearrangements. Genome Res, 14(7), 1394–1403. 10.1101/gr.2289704

Datsenko, K. A., & Wanner, B. L. (2000). One-step inactivation of chromosomal genes in *Escherichia coli* K-12 using PCR products. Proc Natl Acad Sci, 97(12), 6640–6645. 10.1073/pnas.120163297

De la Cruz Barrón, M., Merlin, C., Guilloteau, H., Montargès-Pelletier, E., & Bellanger, X. (2018). Suspended materials in river waters differentially enrich class 1 integron- and IncP-1 plasmid-carrying bacteria in sediments. Front Microbiol, 9(2018). https://www.frontiersin.org/article/10.3389/fmicb.2018.01443

Doumith, M., Ellington, M. J., Livermore, D. M., & Woodford, N. 2009. Molecular mechanisms disrupting porin expression in ertapenem-resistant *Klebsiella* and *Enterobacter* spp. clinical isolates from the UK. J Antimicrob Chemother, 63(4), 659–667. 10.1093/jac/dkp029

Drost, H. (2018). Philentropy: Information Theory and Distance Quantification with R. J Open Source Softw, 3(26), 765. 10.21105/joss.00765

Duan, Y. F., Grogan, P., Walker, V. K., & diCenzo, G. C. (2022). Whole genome sequencing of three mesorhizobia isolated from northern Canada to identify genomic adaptations promoting nodulation in cold climates. Can J Microbiol, 68(11): 661–673. 10.1139/cjm-2022-0102

Evans, B. A., & Amyes, S. G. B. (2014). OXA β-Lactamases. Clin Microbiol Rev, 27(2), 241–263. 10.1128/CMR.00117-13

Galata, V., Fehlmann, T., Backes, C., & Keller, A. (2019). PLSDB: a resource of complete bacterial plasmids. Nucleic Acids Res, 47(D1): D195–D202. 10.1093/nar/gky1050

Gaibani, P., Ambretti, S., Scaltriti, E., Cordovana, M., Berlingeri, A., Pongolini, S., Landini, M. P., & Re, M. C. (2018). A novel IncA plasmid carrying *bla*VIM-1 in a *Kluyvera cryocrescens* strain. J Antimicrob Chemother, 73(11), 3206–3208. 10.1093/jac/dky304

Gillings, M., Boucher, Y., Labbate, M., Holmes, A., Krishnan, S., Holley, M., & Stokes, H. W. (2008). The evolution of class 1 integrons and the rise of antibiotic resistance. J Bacteriol, 190(14), 5095–5100. 10.1128/JB.00152-08

Grant, S. G., Jessee, J., Bloom, F. R., & Hanahan, D. (1990). Differential plasmid rescue from transgenic mouse DNAs into *Escherichia coli* methylation-restriction mutants. Proc Natl Acad Sci, 87(12), 4645–4649.

Guyer, M. S., Reed, R. R., Steitz, J. A., & Low, K. B. (1981). Identification of a sex-factor- affinity site in *E. coli* as gamma delta. Cold Spring Harb Symp Quant Biol, 44(1), 135–140. doi: 10.1101/sqb.1981.045.01.022. PMID: 6271456.

Hawkey, P. M., & Livermore, D. M. (2012). Carbapenem antibiotics for serious infections. BMJ, 344, e3236. 10.1136/bmj.e3236

Hu, Y., Yang, X., Li, J., Lv, N., Liu, F., Wu, J., Lin, I. Y. C., Wu, N., Weimer, B. C., Gao, G. F., Liu, Y., & Zhu, B. (2016). The bacterial mobile resistome transfer network connecting the animal and human microbiomes. Appl Environ Microbiol, 82(22), 6672–6681. 10.1128/AEM.01802-16

Hunt, M., Silva, N. D., Otto, T. D., Parkhill, J., Keane, J. A., & Harris, S. R. (2015). Circlator: Automated circularization of genome assemblies using long sequencing reads. Genome Biol, 16(1), 294. 10.1186/s13059-015-0849-0

Irrgang, A., Tenhagen, B.-A., Pauly, N., Schmoger, S., Kaesbohrer, A., & Hammerl, J. A. (2019). Characterization of VIM-1-producing *E. coli* isolated from a German fattening pig farm by an improved isolation procedure. Front Microbiol, 10(2019). https://www.frontiersin.org/article/10.3389/fmicb.2019.02256

Jacoby, G. A., Mills, D. M., & Chow, N. 2004. Role of β-lactamases and porins in resistance to ertapenem and other β-lactams in *Klebsiella pneumoniae*. Antimicrob Agents Chemother, 48(8), 3203–3206. 10.1128/aac.48.8.3203-3206.2004

Jaffe, A., Chabbert, Y. A., & Semonin, O. (1982). Role of porin proteins OmpF and OmpC in the permeation of beta-lactams. Antimicrob Agents Chemother, 22(6), 942–948.

Jamal, A. J., Mataseje, L. F., Williams, V., Leis, J. A., Tijet, N., Zittermann, S., Melano, R. G., Mulvey, M. R., Katz, K., Allen, V. G., & McGeer, A. J. (2021). Genomic epidemiology of carbapenemase-producing *Enterobacterales* at a hospital system in Toronto, Ontario, Canada, 2007 to 2018. Antimicrob Agents Chemother, 65(8), e00360–21. 10.1128/AAC.00360-21

Johansson, Å., Ekelöf, J., Giske, C. G., & Sundqvist, M. (2014). The detection and verification of carbapenemases using ertapenem and Matrix Assisted Laser Desorption Ionization-Time of Flight. BMC Microbiol, 14, 89. 10.1186/1471-2180-14-89

Katoh, K., Rozewicki, J., & Yamada, K. D. (2019). MAFFT online service: Multiple sequence alignment, interactive sequence choice and visualization. Brief Bioinform, 20(4), 1160–1166. 10.1093/bib/bbx108

Kintses, B., Méhi, O., Ari, E., Számel, M., Györkei, Á., Jangir, P. K., Nagy, I., Pál, F., Fekete, G., Tengölics, R., Nyerges, Á., Likó, I., Bálint, A., Molnár, T., Bálint, B., Vásárhelyi, B. M., Bustamante, M., Papp, B., & Pál, C. (2019). Phylogenetic barriers to horizontal transfer of antimicrobial peptide resistance genes in the human gut microbiota. Nat Microbiol, 4(3), 447–458. 10.1038/s41564-018-0313-5

Knopp, M., & Andersson, D. I. (2015). Amelioration of the fitness costs of antibiotic resistance due to reduced outer membrane permeability by upregulation of alternative porins. Mol Biol Evol, 32(12), 3252–3263. 10.1093/molbev/msv195

Kohler, P. P., Melano, R. G., Patel, S. N., Shafinaz, S., Faheem, A., Coleman, B. L., Green, K., Armstrong, I., Almohri, H., Borgia, S., Borgundvaag, E., Johnstone, J., Katz, K., Lam, F., Muller, M. P., Powis, J., Poutanen, S. M., Richardson, D., Rebbapragada, A., Sarabia, A., Simor, A., McGeer, A., & the Toronto Invasive Bacterial Diseases Network (TIBDN). (2018). Emergence of Carbapenemase-Producing Enterobacteriaceae, South-Central Ontario, Canada. Emerg Infect Dis, 24(9), 1674–1682. 10.3201/eid2409.180164

Kohler, P., Tijet, N., Kim, H. C., Johnstone, J., Edge, T., Patel, S. N., Seah, C., Willey, B., Coleman, B., Green, K., Armstrong, I., Katz, K., Muller, M. P., Powis, J., Poutanen, S. M., Richardson, D., Sarabia, A., Simor, A., McGeer, A., & Melano, R. G. (2020). Dissemination of Verona Integron-encoded Metallo-β-lactamase among clinical and environmental *Enterobacteriaceae* isolates in Ontario, Canada. Sci Rep, 10(1), Article 1. 10.1038/s41598-020-75247-7

Kolmogorov, M., Yuan, J., Lin, Y., & Pevzner, P. A. (2019). Assembly of long, error-prone reads using repeat graphs. Nat Biotechnol, 37(5), 540–546. 10.1038/s41587-019-0072-8

Kovach, M. E., Elzer, P. H., Steven Hill, D., Robertson, G. T., Farris, M. A., Roop, R. M., & Peterson, K. M. (1995). Four new derivatives of the broad-host-range cloning vector pBBR1MCS, carrying different antibiotic-resistance cassettes. Gene, 166(1), 175–176. 10.1016/0378-1119(95)00584-1

Kurtz, S., Phillippy, A., Delcher, A. L., Smoot, M., Shumway, M., Antonescu, C., & Salzberg, S. L. (2004). Versatile and open software for comparing large genomes. Genome Biol, 5(2), R12. 10.1186/gb-2004-5-2-r12

Leavitt, A., Chmelnitsky, I., Colodner, R., Ofek, I., Carmeli, Y., & Navon-Venezia, S. (2009). Ertapenem resistance among extended-spectrum-β-lactamase-producing *Klebsiella pneumoniae* isolates. J Clin Microbiol, 47(4), 969–974. 10.1128/JCM.00651-08

Letunic, I., & Bork, P. (2021). Interactive Tree Of Life (iTOL) v5: an online tool for phylogenetic tree display and annotation. Nucleic Acids Res., 49(W1), W293–W296. 10.1093/nar/gkab301

Li, H. (2018). Minimap2: Pairwise alignment for nucleotide sequences. Bioinformatics, 34(18), 3094–3100. 10.1093/bioinformatics/bty191

Li, H., & Durbin, R. (2009). Fast and accurate short read alignment with Burrows-Wheeler transform. Bioinformatics, 25(14), 1754–1760. 10.1093/bioinformatics/btp324

Li, Y., da Silva, G. C., Li, Y., Rossi, C. C., Fernandez Crespo, R., Williamson, S. M., Langford, P. R., Bazzolli, D. M. S., & Bossé, J. T. (2018). Evidence of illegitimate recombination between two *Pasteurellaceae* plasmids resulting in a novel multi-resistance replicon, pM3362MDR, in *Actinobacillus pleuropneumoniae*. Front Microbiol, 9(2018). https://www.frontiersin.org/article/10.3389/fmicb.2018.02489

Liu, L., Feng, Y., Hu, Y., Kang, M., Xie, Y., & Zong, Z. (2018). *Klebsiella grimontii*, a new species acquired carbapenem resistance. Front Microbiol, 9(2018). 10.3389/fmicb.2018.02170

Mano, Y., Saga, T., Ishii, Y., Yoshizumi, A., Bonomo, R. A., Yamaguchi, K., & Tateda, K. (2015). Molecular analysis of the integrons of metallo-β-lactamase-producing *Pseudomonas aeruginosa* isolates collected by nationwide surveillance programs across Japan. BMC Microbiol, 15(1), 41. 10.1186/s12866-015-0378-8

Matsumura, Y., Peirano, G., Devinney, R., Bradford, P. A., Motyl, M. R., Adams, M. D., Chen, L., Kreiswirth, B., & Pitout, J. D. D. (2017). Genomic epidemiology of global VIM-producing *Enterobacteriaceae*. J Antimicrob Chemother, 72(8), 2249–2258. 10.1093/jac/dkx148

Mattioni Marchetti, V., Bitar, I., Piazza, A., Mercato, A., Fogato, E., Hrabak, J., & Migliavacca, R. (2020). Genomic insight of VIM-harboring IncA plasmid from a clinical ST69 *Escherichia coli* strain in Italy. Microorganisms, 8(8), 1232. 10.3390/microorganisms8081232

Meier-Kolthoff, J. P., Carbasse, J. S., Peinado-Olarte, R. L., & Göker, M. (2022). TYGS and LPSN: A database tandem for fast and reliable genome-based classification and nomenclature of prokaryotes. Nucleic Acids Res, 50(D1), D801–D807. 10.1093/nar/gkab902

Minh, B. Q., Schmidt, H. A., Chernomor, O., Schrempf, D., Woodhams, M. D., von Haeseler, A., & Lanfear, R. (2020). IQ-TREE 2: New models and efficient methods for phylogenetic inference in the genomic era. Mol Bol Evol, 37(5), 1530–1534. 10.1093/molbev/msaa015

Miriagou, V., Cornaglia, G., Edelstein, M., Galani, I., Giske, C. G., Gniadkowski, M., Malamou-Lada, E., Martinez-Martinez, L., Navarro, F., Nordmann, P., Peixe, L., Pournaras, S., Rossolini, G. M., Tsakris, A., Vatopoulos, A., & Cantón, R. (2010). Acquired carbapenemases in Gram-negative bacterial pathogens: Detection and surveillance issues. Clin Microbiol Infect, 16(2), 112–122. 10.1111/j.1469-0691.2009.03116.x

Murray, C. J., Ikuta, K. S., Sharara, F., Swetschinski, L., Aguilar, G. R., Gray, A., Han, C., Bisignano, C., Rao, P., Wool, E., Johnson, S. C., Browne, A. J., Chipeta, M. G., Fell, F., Hackett, S., Haines-Woodhouse, G., Hamadani, B. H. K., Kumaran, E. A. P., McManigal, B., Agarwal, R., Akech, S., Albertson, S., Amuasi, J., Andrews, J., Aravkin, A., Ashley, E., Bailey, F., Baker, S., Basnyat, B., Bekker, A., Bender, R., Bethou, A., Bielicki, J., Boonkasidecha, S., Bukosia, J., Carvalheiro, C., Castañeda-Orjuela, C., Chansamouth, V., Chaurasia, S., Chiurchiù, S., Chowdhury, F., Cook, A. J., Cooper, B., Cressey, T. R., Criollo-Mora, E., Cunningham, M., Darboe, S., Day, N. P. J., De Luca, M., Dokova, K., Dramowski, A., Dunachie, S. J., Eckmanns, T., Eibach, D., Emami, A., Feasey, N., Fisher-Pearson, N., Forrest, K., Garrett, D., Gastmeier, P. Giref, A. Z., Greer, R. C., Gupta, V., Haller, S., Haselbeck, A., Hay, S. I., Holm, M., Hopkins, S., Iregbu, K. C., Jacobs, J., Jarovsky, D., Javanmardi, F., Khorana, M., Kissoon, N., Kobeissi, E., Kostyanev, T., Krapp, F., Krumkamp, R., Kumar, A., Kyu, H. H., Lim, C., Limmathurotsakul, D., Loftus, M. J., Lunn, M., Ma, J., Mturi, N., Munera-Huertas, T., Musicha, P., Mussi-Pinhata, M. M., Nakamura, T., Nanavati, R., Nangia, S., Newton, P., Ngoun, C., Novotney, A., Nwakanma, D., Obiero, C. W., Olivas-Martinez, A., Olliaro, P., Ooko, E., Ortiz-Brizuela, E., Peleg, A. Y., Perrone, C., Plakkal, N., Ponce-de-Leon, A., Raad, M., Ramdin, T., Riddell, A., Roberts, T., Robotham, J. V., Roca, A., Rudd, K. E., Russell, N., Schnall, J., Scott, J. A. G., Shivamallappa, M., Sifuentes-Osornio, J., Steenkeste, N., Stewardson, A. J., Stoeva, T., Tasak, N., Thaiprakong, A., Thwaites, G., Turner, C., Turner, P., Rogier van Doorn, H., Velaphi, S., Vongpradith, A., Vu, H., Walsh, T., Waner, S., Wangrangsimakul, T., Wozniak, T., Zheng, P., Sartorius, B., Lopez, A. D., Stergachis, A., Moore, C., Dolecek, C. & Naghavi, M. (2022). Global burden of bacterial antimicrobial resistance in 2019: A systematic analysis. Lancet, 399(10325), 629–655. 10.1016/S0140-6736(21)02724-0

Néron, B., Littner, E., Haudiquet, M., Perrin, A., Cury, J., & Rocha, E. P. C. (2022). IntegronFinder 2.0: Identification and analysis of integrons across bacteria, with a focus on antibiotic resistance in *Klebsiella*. Microorganisms, 10(4), Article 4. 10.3390/microorganisms10040700

Page, A. J., Cummins, C. A., Hunt, M., Wong, V., K., Reuter, S., Holden, M. T. G., Fookes, M., Falush, D., Keane, J. A., & Parkhill, J. (2015). Roary: rapid large-scale prokaryote pan genome analysis. Bioinformatics, 31(22), 3691–3693. 10.1093/bioinformatics/btv421

Papadimitriou-Olivgeris, M., Bartzavali, C., Lambropoulou, A., Solomou, A., Tsiata, E., Anastassiou, E. D., Fligou, F., Marangos, M., Spiliopoulou, I., & Christofidou, M. (2019). Reversal of carbapenemase-producing *Klebsiella pneumoniae* epidemiology from blaKPC- to blaVIM-harbouring isolates in a Greek ICU after introduction of ceftazidime/avibactam. Journal Antimicrob Chemother, 74(7), 2051–2054. 10.1093/jac/dkz125

Paradis, E., & Schliep, K. (2019). ape 5.0: an environment for modern phylogenetics and evolutionary analyses in R. Bioinformatics, 35(3), 526–528. 10.1093/bioinformatics/bty633

Parks, D.H., Imelfort, M., Skennerton, C.T., Hugenholtz, P., Tyson, G.W. (2015). Assessing the quality of microbial genomes recovered from isolates, single cells, and metagenomes. Genome Res, 25: 1043–1055.

Peirano, G., Lascols, C., Hackel, M., Hoban, D. J., & Pitout, J. D. D. (2014). Molecular epidemiology of *Enterobacteriaceae* that produce VIMs and IMPs from the SMART surveillance program. Diagn Microbiol Infect Dis, 78(3), 277–281. 10.1016/j.diagmicrobio.2013.11.024

Poidevin, M., Sato, M., Altinoglu, I., Delaplace, M., Sato, C., & Yamaichi, Y. (2018). Mutation in ESBL plasmid from *Escherichia coli* O104:H4 leads autoagglutination and enhanced plasmid dissemination. Front Microbiol, 9(2018). https://www.frontiersin.org/article/10.3389/fmicb.2018.00130

Poirel, L., Naas, T., Nicolas, D., Collet, L., Bellais, S., Cavallo, J.-D., & Nordmann, P. (2000). Characterization of VIM-2, a carbapenem-hydrolyzing metallo-β-lactamase and its plasmid- and integron-borne gene from a *Pseudomonas aeruginosa* clinical isolate in France. Antimicrob Agents Chemother, 44(4), 891–897.

Porse, A., Schou, T. S., Munck, C., Ellabaan, M. M. H., & Sommer, M. O. A. (2018). Biochemical mechanisms determine the functional compatibility of heterologous genes. Nat Commun, 9, 522. 10.1038/s41467-018-02944-3

Prjibelski, A., Antipov, D., Meleshko, D., Lapidus, A., & Korobeynikov, A. (2020). Using SPAdes de novo assembler. Curr Protoc Bioinform, 70(1), e102. 10.1002/cpbi.102

R Core Team (2021). R: A language and environment for statistical computing. R Foundation for Statistical Computing, Vienna, Austria. URL https://www.R-project.org/.

Rodriguez-Martinez, J.-M., Nordmann, P., Fortineau, N., & Poirel, L. (2010). VIM-19, a metallo-β-lactamase with increased carbapenemase activity from *Escherichia coli* and *Klebsiella pneumoniae*. Antimicrob Agents Chemother, 54(1), 471–476. 10.1128/AAC.00458-09

Salimraj, R., Hinchliffe, P., Kosmopoulou, M., Tyrrell, J. M., Brem, J., van Berkel, S. S., Verma, A., Owens, R. J., McDonough, M. A., Walsh, T. R., Schofield, C. J., & Spencer, J. (2019). Crystal structures of VIM-1 complexes explain active site heterogeneity in VIM-class metallo-β-lactamases. FEBS J, 286(1), 169–183. 10.1111/febs.14695

Sánchez-Romero, I., Asensio, Á., Oteo, J., Muñoz-Algarra, M., Isidoro, B., Vindel, A., Álvarez-Avello, J., Balandín-Moreno, B., Cuevas, O., Fernández-Romero, S., Azañedo, L., Sáez, D., & Campos, J. (2012). Nosocomial outbreak of VIM-1-producing *Klebsiella pneumoniae* isolates of multilocus sequence Type 15: Molecular basis, clinical risk factors, and outcome. Antimicrob Agents Chemother, 56(1), 420–427. 10.1128/AAC.05036-11

Schulz, G. E. (2002). The structure of bacterial outer membrane proteins. Biochim Biophys Acta, 1565(2), 308–317. 10.1016/s0005-2736(02)00577-1

Seeman, T. (2014). Prokka: rapid prokaryotic genome annotation. Bioinformatics, 30(14), 2068–2069. 10.1093/bioinformatics/btu153

Seemann, T. (2015). snippy: Fast bacterial variant calling from NGS reads [Computer software]. https://github.com/tseemann/snippy

Sievers, F., & Higgins, D. G. (2017). Clustal Omega for making accurate alignments of many protein sequences. Protein Sci., 27, 135–145. 10.1002/pro.3290

Silva, R. F., Mendonça, S. C. M., Carvalho, L. M., Reis, A. M., Gordo, I., Trindade, S., & Dionisio, F. (2011). Pervasive sign epistasis between conjugative plasmids and drug-resistance chromosomal mutations. PLOS Genet, 7(7), e1002181. 10.1371/journal.pgen.1002181

Smillie, C., Garcillán-Barcia, M. P., Francia, M. V., Rocha, E. P. C., & de la Cruz, F. (2010). Mobility of plasmids. Microbiol Mol Biol Rev, 74(3), 434–452. 10.1128/MMBR.00020-10

Socha, R. D., Chen, J., & Tokuriki, N. (2019). The molecular mechanisms underlying hidden phenotypic variation among metallo-β-lactamases. J Mol Biol, 431(6), 1172–1185. 10.1016/j.jmb.2019.01.041

Sorek, R., Zhu, Y., Creevey, C. J., Francino, M. P., Bork, P., & Rubin, E. M. (2007). Genome-wide experimental determination of barriers to horizontal gene transfer. Science, 318(5855), 1449–1452. 10.1126/science.1147112

Streicher, S. L., Shanmugam, K. T., Ausubel, F., Morandi, C. & Goldberg, R. B. (1974). Regulation of nitrogen fixation in *Klebsiella pneumoniae*: evidence for a role of glutamine synthetase as a regulator of nitrogenase synthesis. J Bacteriol, 120(2), 815–821. 10.1128/jb.120.2.815-821.1974

Szabó, D., Silveira, F., Hujer, A. M., Bonomo, R. A., Hujer, K. M., Marsh, J. M., Bethel, C. R., Doi, Y., Deeley, K. & Paterson, D. L. (2006). Outer membrane protein changes and efflux pump expression together may confer resistance to ertapenem in *Enterobacter cloacae*. Antimicrob Agents Chemother, 50(8), 2833–2835. 10.1128/aac.01591-05

Tatusova, T., DiCuccio, M., Badretdin, A., Chetvernin, V., Nawrocki, E. P., Zaslavsky, L., Lomsadze, A., Pruitt, K. D., Borodovsky, M., & Ostell, J. (2016). NCBI prokaryotic genome annotation pipeline. Nucleic Acids Res, 44(14), 6614–6624. 10.1093/nar/gkw569

Tijet, N., Macmullin, G., Lastovetska, O., Vermeiren, C., Wenzel, P., Stacey-Works, T., Low, D. E., Patel, S. N., & Melano, R. G. (2013). Verona Integron–encoded Metallo-β-Lactamase 1 in *Enterobacteria*, Ontario, Canada. Emerg Infect Dis, 19(7), 1156–1158. 10.3201/eid1907.121294

van der Zee, A., Kraak, W. B., Burggraaf, A., Goessens, W. H. F., Pirovano, W., Ossewaarde, J. M., & Tommassen, J. (2018). Spread of carbapenem resistance by transposition and conjugation among *Pseudomonas aeruginosa*. Front Microbiol, 9(2018). https://www.frontiersin.org/article/10.3389/fmicb.2018.02057

Vaser, R., Sović, I., Nagarajan, N., & Šikić, M. (2017). Fast and accurate de novo genome assembly from long uncorrected reads. Genome Res, 27(5), 737–746. 10.1101/gr.214270.116

Ventola, C. L. (2015). The antibiotic resistance crisis. Pharm Ther, 40(4), 277–283.

von Wintersdorff, C. J. H., Penders, J., van Niekerk, J. M., Mills, N. D., Majumder, S., van Alphen, L. B., Savelkoul, P. H. M., & Wolffs, P. F. G. (2016). Dissemination of antimicrobial resistance in microbial ecosystems through horizontal gene transfer. Front Microbiol, 7(2016). https://www.frontiersin.org/article/10.3389/fmicb.2016.00173

Walker, B. J., Abeel, T., Shea, T., Priest, M., Abouelliel, A., Sakthikumar, S., Cuomo, C. A., Zeng, Q., Wortman, J., Young, S. K., & Earl, A. M. (2014). Pilon: An integrated tool for comprehensive microbial variant detection and genome assembly improvement. PloS One, 9(11), e112963. 10.1371/journal.pone.0112963

Wick, R. R., Judd, L. M., Gorrie, C. L., & Holt, K. E. (2017). Unicycler: Resolving bacterial genome assemblies from short and long sequencing reads. PLOS Comput Biol, 13(6), e1005595. 10.1371/journal.pcbi.1005595

Wong, A. (2017). Epistasis and the evolution of antimicrobial resistance. Front Microbiol, 8(2017). https://www.frontiersin.org/articles/10.3389/fmicb.2017.00246

Woodford, D., Dallow, J. W. T., Hill, R. L. R., Palepou, M. I., Pike, R., Ward, M. E., Warner, M. & Livermore, D. M. 2007. Ertapenem resistance among *Klebsiella* and *Enterobacter* submitted in the UK to a reference laboratory. Int J Antimicrob Agents, 29(4), 456–459. 10.1016/j.ijantimicag.2006.11.020

Zeng, L., Zhan, Z., Hu, L., Jiang, X., Zhang, Y., Feng, J., Gao, B., Zhao, Y., Yang, W., Yang, H., Yin, Z., & Zhou, D. (2019). Genetic characterization of a blaVIM-24-carrying IncP-7β plasmid p1160-VIM and a blaVIM-4-harboring integrative and conjugative element Tn6413 from clinical *Pseudomonas aeruginosa*. Front Microbiol, 10, 213. 10.3389/fmicb.2019.00213

