## Supplementary Materials for "Variations in carbapenem resistance associated with the VIM-1 metallo-β-lactamase across the *Enterobacterales*"

**Supplementary Material for**
Variations in carbapenem resistance associated with the VIM-1 metallo-β-lactamase across the order *Enterobacterales*

Mia Rondinelli, Sabhjeet Kaur, Owen A. Ledwell, Henry Wong, Prameet M. Sheth, George C. diCenzo

George C. diCenzo


**This PDF file includes**

Tables S1 to S2 (pages 2-4)
Figures S1 to S2 (pages 5-6)

**Table S1.** Oligonucleotides used in this study.

| Primer name | Sequence | Application |
|---|---|---|
| MR001 | 5'-CCC CAA GCT TCT AGG TTA TGG AGC AGC AAC GAT GTT AC | Forward primer for amplifying *vim-1* of *E. coli* F5446 (encapsulates *attC* site on 3' end) |
| MR003 | 5'-GCG CGG ATC CGC AAA AGT CCC GCT CCA AGC | Reverse primer for amplifying *vim-1* of *E. coli* F5446 |
| SK003 | 5'-ACT CCT ACG GGA GGC AGC AGT | Forward primer for qPCR of 16S rRNA genes. |
| SK004 | 5'-TAT TAC CGC GGC TGC TGG C | Reverse primer for qPCR of 16S rRNA genes. |
| GD018 | 5'-ATG GTC TCA TTG TCC GTG ATG | Forward primer for screening VIM in *Enterobacteriaceae* clinical isolates and for qPCR. |
| GD019 | 5'-GTC ATG AAA GTG CGT GGA GA | Reverse primer for screening VIM in *Enterobacteriaceae* clinical isolates and for qPCR. |

Restriction sites added to the 5' end of primers to facilitate clonings are underlined.

**Table S2.** Summary of resistance genes identified in the clinical isolates using the Resistance Gene Identifier (RGI) of the Comprehensive Antibiotic Resistance Database (CARD).

| Gene name | AMR gene family | Resistance mechanism | F5446 | S2568 | F48994 | H17629 | H70375 | T64870 |
|---|---|---|---|---|---|---|---|---|
| *AAC(6')-Ib-cr5* | AAC(6') aminoglycoside acetyltransferase | Antibiotic inactivation | ✓ | | ✓ | | ✓ | |
| *AAC(6')-Ib'* | AAC(6') aminoglycoside acetyltransferase | Antibiotic inactivation | | | ✓ | ✓ | | |
| *AAC(6')-Ib4* | AAC(6') aminoglycoside acetyltransferase | Antibiotic inactivation | ✓ | ✓ | ✓ | ✓ | ✓ | ✓ |
| *ACT-24* | ACT beta-lactamase | Antibiotic inactivation | | | ✓ | | | |
| *aadA3* | ANT(3") nucleotidyltransferase | Antibiotic inactivation | | | | | ✓ | |
| *aadA5* | ANT(3") nucleotidyltransferase | Antibiotic inactivation | ✓ | ✓ | | | | |
| *aph(3')-XV* | APH(3') aminoglycoside O-phosphotransferase | Antibiotic inactivation | ✓ | ✓ | | | | |
| *aphA15* | APH(3') aminoglycoside phosphorylase | Antibiotic inactivation | ✓ | ✓ | | | | |
| *LptD* | ATP-binding cassette (ABC) antibiotic efflux pump | Antibiotic efflux | | | | | | ✓ |
| *msbA* | ATP-binding cassette (ABC) antibiotic efflux pump | Antibiotic efflux | ✓ | ✓ | | | | |
| *CTX-M-15* | CTX-M beta-lactamase | Antibiotic inactivation | ✓ | ✓ | ✓ | | ✓ | |
| *LAP-2* | LAP beta-lactamase | Antibiotic inactivation | | | ✓ | | | |
| *mphA* | Macrolide phosphotransferase (MPH) | Antibiotic inactivation | ✓ | ✓ | ✓ | | | ✓ |
| *emrB* | Major facilitator superfamily (MFS) antibiotic efflux pump | Antibiotic efflux | ✓ | ✓ | | | | |
| *emrR* | Major facilitator superfamily (MFS) antibiotic efflux pump | Antibiotic efflux | ✓ | ✓ | | | | |
| *emrY* | Major facilitator superfamily (MFS) antibiotic efflux pump | Antibiotic efflux | ✓ | ✓ | | | | |
| *K. pneumoniae KpnE* | Major facilitator superfamily (MFS) antibiotic efflux pump | Antibiotic efflux | | | | | | ✓ |
| *K. pneumoniae KpnF* | Major facilitator superfamily (MFS) antibiotic efflux pump | Antibiotic efflux | | | | | | ✓ |
| *mdtG* | Major facilitator superfamily (MFS) antibiotic efflux pump | Antibiotic efflux | ✓ | ✓ | | | | |
| *mdtH* | Major facilitator superfamily (MFS) antibiotic efflux pump | Antibiotic efflux | ✓ | ✓ | | | | |
| *qacEdelta1* | Major facilitator superfamily (MFS) antibiotic efflux pump | Antibiotic efflux | ✓ | ✓ | ✓ | ✓ | ✓ | ✓ |

*Continued on next page*

| Gene name | AMR gene family | Resistance mechanism | F5446 | S2568 | F48994 | H17629 | H70375 | T64870 |
|---|---|---|---|---|---|---|---|---|
| *evgA* | Major facilitator superfamily (MFS) antibiotic efflux pump, resistance-nodulation-cell division (RND) antibiotic efflux pump | Antibiotic efflux | ✓ | ✓ | | | | |
| *H-NS* | Major facilitator superfamily (MFS) antibiotic efflux pump, resistance-nodulation-cell division (RND) antibiotic efflux pump | Antibiotic efflux | ✓ | ✓ | | | | |
| *OXA-1* | OXA beta-lactamase | Antibiotic inactivation | ✓ | | ✓ | | ✓ | |
| *OXA-9* | OXA beta-lactamase | Antibiotic target replacement | | | ✓ | | | |
| *QnrB1* | Quinolone resistance protein (qnr) | Antibiotic target protection | | | | | ✓ | |
| *QnrS1* | Quinolone resistance protein (qnr) | Antibiotic target protection | | | ✓ | ✓ | ✓ | ✓ |
| *baeR* | Resistance-nodulation-cell division (RND) antibiotic efflux pump | Antibiotic efflux | ✓ | ✓ | | | | |
| *oqxA* | Resistance-nodulation-cell division (RND) antibiotic efflux pump | Antibiotic efflux | | | | | | ✓ |
| *marA* | Resistance-nodulation-cell division (RND) antibiotic efflux pump, General Bacterial Porin with reduced permeability to beta-lactams | Antibiotic efflux, reduced permeability to antibiotic | ✓ | ✓ | | | | |
| *SHV-12* | SHV beta-lactamase | Antibiotic inactivation | ✓ | ✓ | | | | |
| *SHV-28* | SHV beta-lactamase | Antibiotic inactivation | | | | | | ✓ |
| *sul1* | Sulfonamide resistant sul | Antibiotic target replacement | ✓ | ✓ | ✓ | ✓ | ✓ | ✓ |
| *sul2* | Sulfonamide resistant sul | Antibiotic target replacement | | | | | ✓ | |
| *TEM-1* | TEM beta-lactamase | Antibiotic inactivation | | | ✓ | | ✓ | |
| *dfrA14* | Trimethoprim resistant dihydrofolate reductase dfr | Antibiotic target replacement | ✓ | ✓ | ✓ | ✓ | ✓ | ✓ |
| *dfrA15* | Trimethoprim resistant dihydrofolate reductase dfr | Antibiotic target replacement | | | ✓ | | | |
| *dfrA17* | Trimethoprim resistant dihydrofolate reductase dfr | Antibiotic target replacement | ✓ | ✓ | | | | |
| *VIM-1* | VIM beta-lactamase | Antibiotic inactivation | ✓ | ✓ | ✓ | ✓ | ✓ | ✓ |

F5446 – *E. coli*; S2568 – *E. coli*; F48994 – *Enterobacter hormaechei*; H17629 – *E. hormaechei*; H70375 – *E. hormaechei*; T64870 – *Klebsiella pneumoniae*.

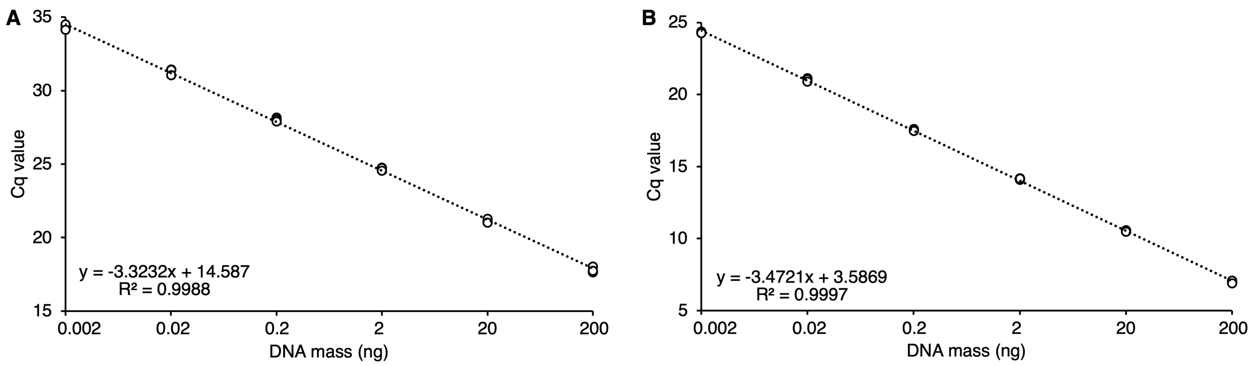


**Figure S1. qPCR standard curves.** Standard curves are shown for the primers (**A**) GD018 and GD019 targeting *vim-1*, and (**B**) SK003 and SK004 targeting the 16S rRNA gene. Technical triplicates were performed for each tested cDNA concentration and are plotted separately. The Y-axis represents the total mass of cDNA added to each reaction. The dashed line shows the linear line-of-best-fit. Based on the slope, the amplification efficiency for the *vim-1* primers is 99.9% while the amplification efficiency for the 16S rRNA gene primers is 94.1%.

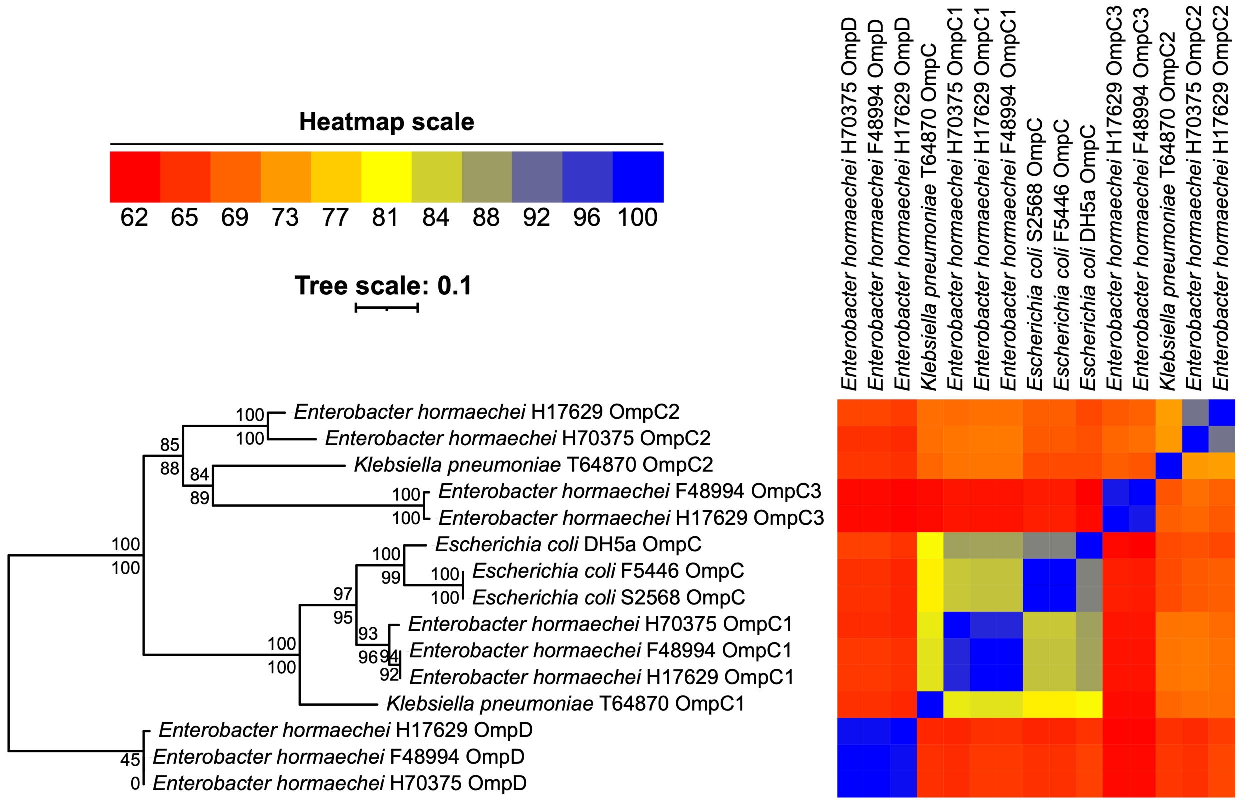


**Figure S2. Phylogeny of OmpC and OmpD of several strains from the order *Enterobacterales*.** OmpC and OmpD proteins were extracted from the proteomes of *Escherchia coli* DH5α and the six clinical isolates described in this study, based on their annotated gene names. On the left is an unrooted maximum likelihood phylogeny showing the evolutionary relationships between the proteins. The numbers on the nodes indicate the ultra-fast bootstrap values (top numbers) and the SH-aLRT support values (bottom numbers), both calculated from 1000 replicates. The scale bar represents the average number of amino acid substitutions per site. On the right is a heatmap showing the percent identity between each pair of proteins, as calculated by Clustal Omega, with colour representing percent identity as shown by the heatmap scale.
